# Hemispheric Fingerprinting: Identifying left and right hemispheres using functional connectivity

**DOI:** 10.64898/2025.12.12.694044

**Authors:** Trevor K. M. Day, Peter E. Turkeltaub, Elissa L. Newport, Andrew T. DeMarco

**Affiliations:** Center for Brain Plasticity and Recovery, Georgetown University, Washington, D.C., U.S.A; MedStar National Rehabilitation Hospital, Washington, D.C., U.S.A

**Keywords:** hemispheric organization, handedness, right-handedness, left-handedness, connectome

## Abstract

Many studies have analyzed what organizational features distinguish the left and right hemispheres of the human brain, with differences typically being found in language areas and in fine motor control (i.e., handedness). In this analysis, we test whether supervised learning can categorize (“fingerprint”) an unseen hemisphere as right or left based on functional connectivity. Using data from the Human Connectome Project, we find success to be extremely high (accuracies > .90) in models trained on right-handed participants (Edinburgh Handedness Inventory [EHI] > 0) and in models trained on left-handed participants (EHI ≤ 0). In a second analysis, we test whether the same can be done to identify handedness alongside hemisphere left/right sidedness. While individuals’ hemispheres are less distinct the more left-handed they are, hemiconnectomes cannot be reliably classified as belonging to a left- or right-handed person. Our approach can inform developmental and post-injury work on hemispheric organization and reorganization.

## 1 Introduction

At first glance, the two hemispheres of the human brain seem to be mirror images of each other. The gross anatomical features and gyral patterns are symmetrical to the naked eye, and the cortex is organized into about a dozen functional networks that are largely symmetric (Hermosillo et al., 2024; Ji et al., 2019; Kong et al., 2025; Power et al., 2011; Yeo et al., 2011).

However, some aspects of function are lateralized, where one hemisphere is specialized to support the function. Most prominently, the language network (LN) is left-lateralized in adults (Lipkin et al., 2022; Poeppel, 2003; Skeide & Friederici, 2016) and becomes more lateralized over development (Olulade et al., 2020; Szaflarski, Holland, et al., 2006, 2006). Left hemisphere (LH) damage results in aphasia — impairment of productive and receptive language — much more frequently than damage to the right hemisphere (RH) (Dewarrat et al., 2009; Turkeltaub, 2019). There are also structural asymmetries in the language network, with the arcuate fasciculus being the most consistently lateralized white-matter tract (Glasser & Rilling, 2008; Lebel & Beaulieu, 2009). Strong leftward laterality is considered to be a bioindicator of healthy development, and damage to the LN has also been hypothesized to be a root cause of schizophrenia (Crow, 2000), in which laterality is reduced (Agcaoglu et al., 2018; Son et al., 2017; Spironelli et al., 2023), as is the case in other conditions, including language and literacy impairments (Bishop, 2013), ADHD (Hale et al., 2005, 2007), and ASD (Floris et al., 2020; Kleinhans et al., 2008), among others.

In contrast, the RH preferentially supports visuospatial processing, including face processing (Kanwisher & Yovel, 2006; Lochy et al., 2019) and attention. The ventral attention network (VAN) is lateralized to the RH (Corbetta et al., 2008; Corbetta & Shulman, 2002) and forms an approximate “mirror” of the LH LN (Bernard et al., 2020; Day, 2024; Ji et al., 2019; Lee et al., 2012; Strike et al., 2023). Unilateral spatial neglect typically follows RH, but not LH injury (Corbetta & Shulman, 2011; Ocklenburg & Güntürkün, 2024; Stone et al., 1993). Additionally, the dorsal attention network (DAN) is also right-lateralized (Wang et al., 2014) and the RH halves of these RSNs are more variable compared to the LH (Perez et al., 2023; Seitzman et al., 2019), suggesting differences in broad connectivity patterns between the two hemispheres. Finally, the frontoparietal network (FPN) connects preferentially to different networks in each hemisphere (Wang et al., 2014).

### 1.1 Handedness

Lateralization of the LN varies across the population, most notably in association with handedness. While the vast majority of right-handed (“dextrals”) are left-lateralized for language (94 – 96%), the rate of left-lateralization is somewhat lower (73 – 78%) in left-handed individuals (“sinistrals”) (Knecht et al., 2000; Szaflarski et al., 2002). This is closely related to the higher cortical heterogeneity in sinistrals, to the point where they are systematically excluded from most neuroscience work, see Willems and colleagues (2014).

Despite this exclusion, sinistrals are not perfectly reversed cortically, and there is conflicting evidence on structural differences in the brain between sinistrals and dextrals, but the largest study to date (*n* dextral *= 28,802; n* sinistral = *3,062)* detected small differences in surface area and thickness between the groups, especially in language and executive function regions (Sha, Pepe, et al., 2021), and there are differences in structural network composition between sinistrals and dextrals (Caeyenberghs & Leemans, 2014).

The exclusion of sinistrals from neuroscience work is predicated on the understanding that their cortices are organized sufficiently distinctly from dextrals to the point that including them would result in greater variance in estimates of brain organization. Thus, we sought to discover whether connectomes could be reliably classified as belonging to sinistrals.

### 1.2 Classification

Prior work has identified differences in function and organization between the two hemispheres; however, it is unknown whether these differences are sufficient to identify a hemisphere with a set of properties as left or right. We use classification algorithms to predict whether an unlabeled connectivity matrix belongs to a left or right hemisphere. While the ground truth is known, classification success has been used to demonstrate the existence of neural signals related to the classification target. For example, connectivity has previously been used to identify individuals out of a group (Finn et al., 2015; Hannum et al., 2023; Kardan et al., 2022; Miranda-Dominguez et al., 2014), predict continuous variables, e.g., age in young children (Kardan et al., 2022), and discrete variables, like sex (Weber et al., 2022; C. Zhang et al., 2018).

Developing a classifier for hemispheres might reveal which within-hemisphere connections are related to hemispheric specialization. Furthermore, such a classifier would permit future work to ask how a developing hemisphere becomes more adult-like in its organization over age, or how an hemisphere contralateral to an injured one is organized after a stroke, e.g. is it more like a typical right or left hemisphere? Here, we use supervised learning to test whether the “sidedness” of a hemisphere is identifiable, i.e., to determine if it is a right hemisphere or a left hemisphere.

### 1.3 Current analysis

We use supervised learning, specifically linear discriminant analysis (LDA) to attempt to classify hemisphere sidedness from connectivity. Specifically, we analyze resting-state fMRI (rsfMRI) data, which is generalizable and collected in many modern studies.

This paper is structured in two major analyses. The first attempts to classify within-hemisphere connectomes (“hemiconnectomes”) as belonging to a right or left hemisphere in separate groups of dextral and sinistral individuals. We investigate the connections that drive the classification and to which network(s) they belong. Finally, the second then attempts to classify hemiconnectomes and complete connectomes as belonging to a sinistral or dextral individual.

We hypothesized that the models would be able to identify hemisphere sidedness, at least in dextrals. We anticipated that hemispheric classification would be more accurate than handedness classification. Finally, we anticipated that connections between auditory and language areas would drive the differences between the hemispheres. Taken together, our goal was to develop an approach that could reliably classify connectomes as belonging to a left or right hemisphere or a left- or right-handed person. With successful classification, the underlying feature weights would help us understand which aspects of cortical organization are different between the two hemispheres or the two handedness groups. A successful classifier could be thought of as a “hemispheric fingerprinter” that could then be used to investigate changes in hemispheric specialization over the lifespan or differences in hemispheric specialization associated with psychiatric or neurological disorders.

## 2 Analysis 1: Hemispheric classification in single-handedness groups

### 2.1 Methods

Our goal is to predict hemispheric sidedness from within-hemisphere connectomes, “hemiconnectomes.” Because auditory and language processing is strongly lateralized, we also examine connectomic features calculated from scans collected while participants performed a language task.

We focus on linear discriminant analysis (LDA) because it provides interpretable parameters, namely, (1) for each training feature, a weight indicating its importance to the classification, and (2) for each observation, a linear discriminant score representing that observation’s position in *n*-dimensional space (for small *n*), based on the weighting of its features. We also replicate some analyses with other models that also show high performance on many-to-one classification problems, neural nets (NN) and support vector classification (SVC), c.f. Hannum et al. (2023). All analyses use publicly available, anonymized data from the Human Connectome Project (HCP) and were reviewed by the Georgetown University Institutional Review Board.

#### 2.1.1 Participants

We used data from 971 HCP (Van Essen et al., 2013) participants with resting-state or language task data. From the demographic variables, we use participant age in whole years, gender (values: male or female), and handedness. The average age of the sample was 28.8 (3.7) y, range: 22 – 37 y, and the entire sample was 54% female.

Handedness was quantified with the Edinburgh Handedness Inventory (Oldfield, 1971), a widely used metric of handedness. EHI scores vary between 100 (strongly dextral) and −100 (strongly sinistral), where absolute value can be taken as an index of handedness strength, regardless of dextrality. The average EHI for the dextral group was 78.4 (19.4) and for the sinistral group, −54.8 (28.2).

Our methods seek to separate within-hemisphere connectivity and connectivity related to handedness by modeling a four-way distinction. However, it is important to note here that the discrete handedness groups are created out of a continuous measurement, the EHI, which does not show discontinuities around EHI = 0. We chose an EHI cutoff of 0, but other cutoffs between dextrals and “non-dextrals” have been proposed.

#### 2.1.2 Fold creation

Participants with either resting state or language task data (total *n = 971*) were divided into cross-validation folds balanced on gender, age, and handedness, see Supplemental Material 1. Folds were created based on the complete *n = 971*, but 935 participants had resting-state data and 963 had language task data, and 927 had both task states.

The dataset was separated into five folds based on age, sex, and handedness status, see SM 1.1 – 1.3. Each analysis was repeated for each fold, i.e., test fold A was tested against models trained on folds B-E, such that each individual is trained on four times, but only tested once. Individuals from the same family were kept in the same fold, and both hemispheres from each individual were always kept together. From these folds, we first create dextral (EHI > 0) and sinistral (EHI ≤ 0) groups for Analysis 1.

In the dextral group, there were 875 total individuals with resting-state and/or language task data. This group was 45% male, mean age: 28.8 y. The sinistral group, n = 96, was 53% male, mean age: 28.8 y. As comparisons for the smaller sinestral group, we created two sets of similarly sized dextral groups, one with random selection (dextral-random, fold sizes: 12 – 27) and one where the dextrals were matched on gender, age, and absolute EHI value to account for differences in absolute EHI variance (dextral-matched, fold sizes: 24 – 54).

#### 2.1.3 Image acquisition

Minimally preprocessed data was accessed from the HCP repository, ConnectomeDB (db.connectome.org). The acquisition parameters and minimal preprocessing have been described previously (Glasser et al., 2013; Van Essen et al., 2013). We retrieved the resting state (RS) and language task (LgT) data, both of which were acquired with a TR of 0.7 s, and 2 mm isotropic voxels. For each participant, two 15 min RS scans (total: 30 min) were acquired as well as two 3:47 LgT scans (total: 7:34). Because language processing is known to be strongly lateralized, we also analyze the BOLD data collected during the LgT, with the goal of evaluating whether strongly correlated evoked connectivity could help classifier performance. The HCP LgT interleaves auditory story blocks (5 – 9 sentences; average duration: 29.8 (SD: 14.5) sec) and math blocks (duration varied based on difficulty). After each block, the participant responded to a two-alternative, forced-choice question about the topic of the story or the answer to the math problem (Barch et al., 2013; Binder et al., 2011).

#### 2.1.4 Atlas selection

We use the Glasser atlas (Glasser et al., 2016), which has 360 parcels, 180 in each hemisphere, resulting in 64,440 unique parcel-to-parcel connections. These connections comprise the full connectome. We divide the full connectome into two hemiconnectomes (HCs) consisting of only within-hemisphere connections, each *n = 16,110.* All 180 parcels appear in both hemispheres.

Because we were interested in differences by hemisphere, and the atlas parcels weren’t exactly symmetric, we conducted several robustness analyses. Specifically, we created two versions of the Glasser atlas where the LH parcels were projected onto the RH (Glasser-LL) and vice versa, where the RH parcels were projected onto the LH (Glasser-RR), which are available on GitHub (Day, 2026/2026).

To aid in interpretation, we use RSNs created from the Glasser atlas and HCP-YA data. Each parcel is assigned to one RSN based on the Cole-Anticevic RSNs (Cole & Ito, 2018; Ji et al., 2019). The Cole-Anticevic networks are not perfectly symmetric, and when homologous parcels were assigned to different networks, the LH assignment was selected to maximize the size of the LN, *n = 14*. Five parcels were assigned to the LN in the LH, but different networks in the RH^1^, see Supplemental Material 2 for more detail. There were no parcels that were assigned to the LN in the RH, but not in the LH.

Of the 12 Cole-Anticevic networks, we merge their two visual networks, since one has only three parcels. The small posterior and ventral multimodal (PMN, VMN) and orbito-affective networks (OAN) are generally ignored (all with three or fewer parcels). The auditory network (AudN) is also small, but kept, due to its relationship with the lateralized LN, see Table A1.M1.

**Table A1.M1:** Cole-Anticevic RSNs, by number of parcels in each hemisphere, and in sum. Abbreviations are given. *: small networks that are not analyzed further.

| Network | Abbrev. | LH | RH | Total |
| --- | --- | --- | --- | --- |
| Auditory | AudN | 8 | 7 | 15 |
| Cingulo-opercular | CON | 27 | 29 | 56 |
| Default | DMN | 40 | 37 | 77 |
| Dorsal attention | DAN | 12 | 11 | 23 |
| Frontoparietal | FPN | 22 | 28 | 50 |
| Language | LN | 14 | 9 | 23 |
| * *Orbito-affective* | *OAN* | 3 | 3 | 6 |
| * *Posterior multimodal* | *PMN* | 3 | 4 | 7 |
| Somatomotor | SMN | 19 | 20 | 39 |
| * *Ventral multimodal* | *VMN* | 2 | 2 | 4 |
| Visual | VisN | 3 | 3 | 6 |
| Visual 2 |  | 27 | 27 | 54 |

#### 2.1.5 Preprocessing

Data were further preprocessed using the eXtensible Connectivity Pipeline-DCAN, (XCP-D) ver. 0.10.6 (Ciric et al., 2018; Mehta et al., 2024; Salo et al., 2025; Satterthwaite et al., 2013), see Supplemental Material 3 for the processing “boilerplate” from XCP-D.

Before model training, Pearson’s *r* values were Fisher-Z transformed. Additionally, to remove the effects of baseline connectivity differences between the hemispheres, the Fisher-Z values were mean-centered and normalized within each hemisphere.

We also calculated hemiconnectomes from the language task data (LgHC), processing it identically, ignoring the task structure. We did not regress out the task, as we anticipated that the increased connectivity in language areas due to a task that creates lateralized activation provides increased information for the models to identify hemisphere sidedness by increasing the magnitude of lateralized connections.

#### 2.1.6 Model assessment

We need a single value to assess model performance, both between models and relative to random chance. Rather than raw classification accuracy, we use Matthew’s correlation coefficient (MCC) to characterize model performance. MCC, like accuracy, sensitivity, or specificity, operates on confusion matrices, but factors in all cells and falls on [-1, 1]. MCC is scaled such that random assignment in one class results in a score of MCC *= 0,* regardless of overall classification success (Chicco & Jurman, 2023; Rainio et al., 2024; Yilmaz & Demirhan, 2023). We use MCC rather than accuracy, because the latter can be misleading in analyses with imbalanced classes, as in Analysis 2. For example, if the model assigns all participants to be dextral, even if the 10% of the sample was sinistral, accuracy would be 90%, which is misleading, since the model is wrong about half of the data types.

There is no analytic standard for calculating *p* values for MCC. However, the relationship is not linear, and simulations suggest that for the present analysis, MCCs of 0.4 and 0.6 are approximately equivalent to 80% and 90% accuracy in each class, see Supplemental Material 4. For all analyses, we adopt these benchmarks of MCC = 0.4 as “good” and 0.6 as “excellent.”

For each set of model results, we bootstrap a 95% confidence interval for that model’s MCC by randomly re-sampling the ground truth and predicted labels with replacement to generate a distribution (Ojala & Garriga, 2009) of MCC, *n = 1,000* repetitions. The mean of this distribution is taken as the MCC value, and the 2.5th and 97.5th percentile of this distribution is used as the 95% confidence interval for the MCC value (Hesterberg, 2014).

#### 2.1.7 Feature selection

The number of features (16,110) exceeds the number of observations (about 2,000 hemispheres total), raising the question of whether fewer features, balanced against the number of observations, can be used to perform sidedness classification. We use the feature selection toolkit from scikit-learn (ver. 1.9.0; Pedregosa et al., 2011), namely SelectKBest(), a supervised selector that returns the best *k* features based on an *F* classifier.

Two common rules of thumb for selecting *k* are (1) 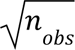 (Hua et al., 2005) or (2) 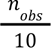 (Steyerberg et al., 2000). We include both of these, as well as *k* taking the value of 2*^x^* from x = 0 to x = 13 (i.e., *k = 8,192*). We repeat this process for both the dextral and sinistral samples.

#### 2.1.8 Identification of important features by network

In order to identify what kind of networks are important, we follow an “enrichment” approach (Eggebrecht et al., 2017). Each connection is a feature in the LDA, and can either be within-network or between-networks.

First, we identify statistically significant features using permutation testing. We randomly shuffle the labels on the input data and re-run the models on this data, creating null models. We repeat this process 10,000 times and extract the feature scalings for the null models, calculate the empirical significance level for the feature scaling from the real data against these repetitions, using the get_p_value() function from the R package infer (ver. 1.1.0; Couch et al., 2021), i.e., a two-sided rank proportion test.

Following this test, connections are marked as significant if the uncorrected *p*-value is less than 0.01. Then, we permute a null distribution of connections across the brain by randomly shuffling those significant values across the connections. Our connectome matrices are symmetric, thus our permutation procedure also maintains the symmetry of the matrix. We then summarize the number of significant connections for all networks and all pairwise networks, and again use get_p_value() to establish if there are more significant connections (regardless of which hemisphere has the stronger connection) in that network pair than expected by chance.

Each pair of networks (or set of within-network connections receives a single *p* value that is later corrected for multiple comparisons. We use false-discovery rate (FDR) correction for 36 multiple comparisons. Simulated *p*-values of 0.0 are replaced with values of .0003 for correction, based on information from the infer manual^2^, which states that the minimum simulated *p*-value scales with *3 / repetitions*, which was 10,000.

Previously, we removed the small networks from the permutation tests, although their connections were included in the training data, so we perform a robustness analysis that re-includes the small, excluded networks (Supplemental Material 7.1). Secondly, because some parcels are assigned to different networks in each hemisphere, this raises the concern that any identified differences between hemispheres are due changing RH network labels to the LH label. In a second robustness test, we exclude entirely the ROIs that belong to different networks in each hemisphere due to atlas asymmetry (Supplemental Material 7.2).

#### 2.1.9 Hemisphere scores

An LDA provides *n - 1* linear discriminants (LD), where *n* is the number of classes. Thus, predicting two hemisphere types results in a single LD, i.e. LD1. Each hemisphere therefore receives an LD1 value on a “hemisphere axis.” We investigated the LD1 values — the hemisphere “fingerprint” — for each of the hemispheres for new models trained on the dextral and sinistral groups.

Because we are no longer testing model reliability, and instead examining model features, we train maximal models on *all* dextral and *all* sinistral participants separately, since model features like LD1 values cannot easily be compared between the five-fold models. This analysis uses all 16,110 features; reductions in the number of features reduces variability in LD1 scores.

The magnitude of the scores varies with both the number of observations and the number of features, and cannot be directly compared between the model trained on dextrals and the model trained on sinistrals. Therefore, we train a third model including both dextrals and sinistrals, and calculate a hemisphere axis “distance” score, where greater “distance” represents hemispheres that are more dissimilar to each other.

#### 2.1.10 Modeling

The primary set of analyses use the discriminant_analysis. LinearDiscriminantAnalysis() function from the Python package scikit-learn (ver. 1.9.0; Pedregosa et al., 2011) in Python 3.13. Ancillary analyses use the neural_network.MLPClassifer() and svm.SVC() functions from the same package, see Supplemental Material 4.

### 2.2 Results

#### 2.2.1 Model performance

The first analysis asks whether hemiconnectomes can be reliably classified as left or right based on a number of choices of training data. The LDA perfectly classified all 1,870 hemispheres, MCC = 1.0 [1.0, 1.0]. Using only one hemisphere from each participant, only four hemispheres were misclassified (2 LH; 2 RH), overall MCC = .991 [.983, 1.0].

Using the language task data, performance was similarly high in both the both-hemispheres training data (MCC = .994 [.989, .999]) and one-hemispheres training data (MCC = .973 [.958, .988]). We replicate these results with NN and SVC, see Supplemental Material 5.1, where the worst-performing model (language task, one hemisphere, NN) still had a near-ceiling MCC of .962.

**Figure A1.R1:**
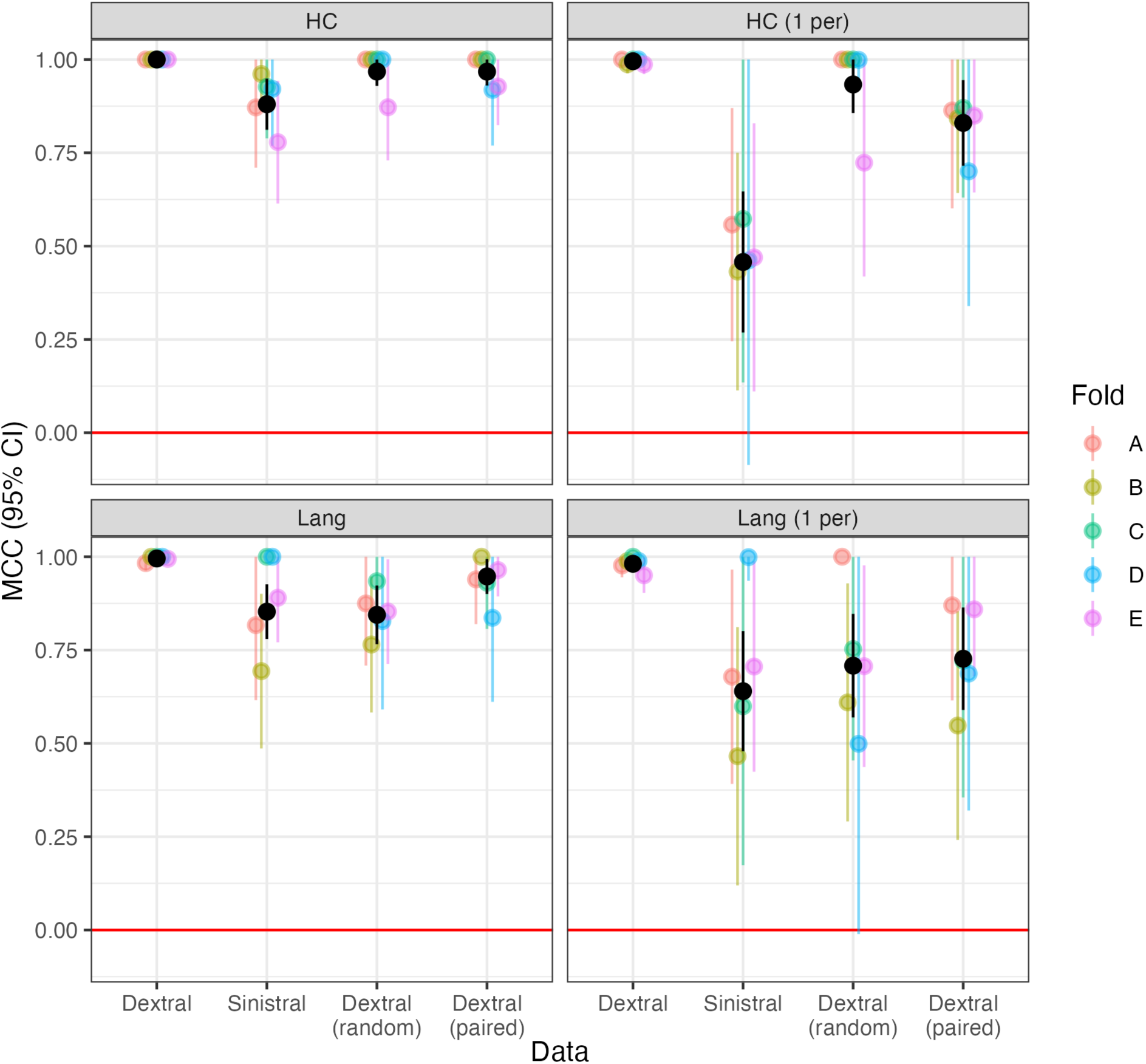
MCC and bootstrapped 95% CI for hemisphere identification by input data type (HC: resting-state; Lang: language task) and by handedness (RHD: dextral; LHD: sinistral) and including the subsampled dextrals (“random” and “paired”). The red line at MCC = 0 represents chance assignment. Models trained and tested on only one hemisphere per participant are included in the right-hand column (“1 per”).

Based on these performance values, we estimated an ANOVA regressing MCC on fold, input data (HC or LT), one-hemisphere inclusion status, and handedness (sinistral, dextral, or the two dextral comparison groups) to examine whether there was a consistent effect of each change to the input data. We found no effect of fold on MCC, *F(4) = 0.942, p = .45;* no difference between task states, *F(1) = 1.7, p = .19*; a reduction in performance when only one hemisphere is included per participant, *F(1) = 26.8, p < .001*; and a difference between the four handedness groups, *F(3) = 15.0, p < .001*.

Consequently, we ran a Tukey post-hoc test on the handedness groups, see Table A1.R1, which shows that the sinistral group performed worse than the dextral group, *β = -.255, p_adj_ < .001.* This is true even in comparison with the groups matched in size, *β = -.139, p_adj_ = .003,* and matched both in size and demographics, *β = -.130, p_adj_ = .006.* The two matched groups underperformed the complete dextral group, but there was no difference between dextral-random and dextral-matched. This suggests that models trained on sinistrals only perform worse, but not due to reductions in training data size or in changes in demographics.

**Table A1.R1:** Paired differences tested between the handedness groups.

| Comparison | Difference (95% CI) | p |
| --- | --- | --- |
| Sinistral - Dextral | -.255 [-.356, -.155] | .000 |
| Sinistral - Dextral (paired) | -.130 [-.230, -.029] | .006 |
| Sinistral - Dextral (random) | -.139 [-.239, -.039] | .003 |
| Dextral (paired) - Dextral | -.126 [-.226, -.025] | .008 |
| Dextral (random) - Dextral | -.116 [-.217, -.016] | .017 |
| Dextral (random) - Dextral (paired) | .009 [-.091, .110] | .995 |

#### 2.2.2 Feature selection

There are many more features (16,110) than observations (1,870 at the largest). Thus, we examine whether the classifier is successful when the number of features is smaller than the number of observations, to avoid over-fitting. Because the sinistral sample is different from the dextral sample, we performed *k*-best feature selection on each separately. Hemisphere identification was MCC > .64 with only one feature, and stably exceeded MCC > .90 for both dextrals and sinistrals with 256 features.

**Figure A1.R2:**
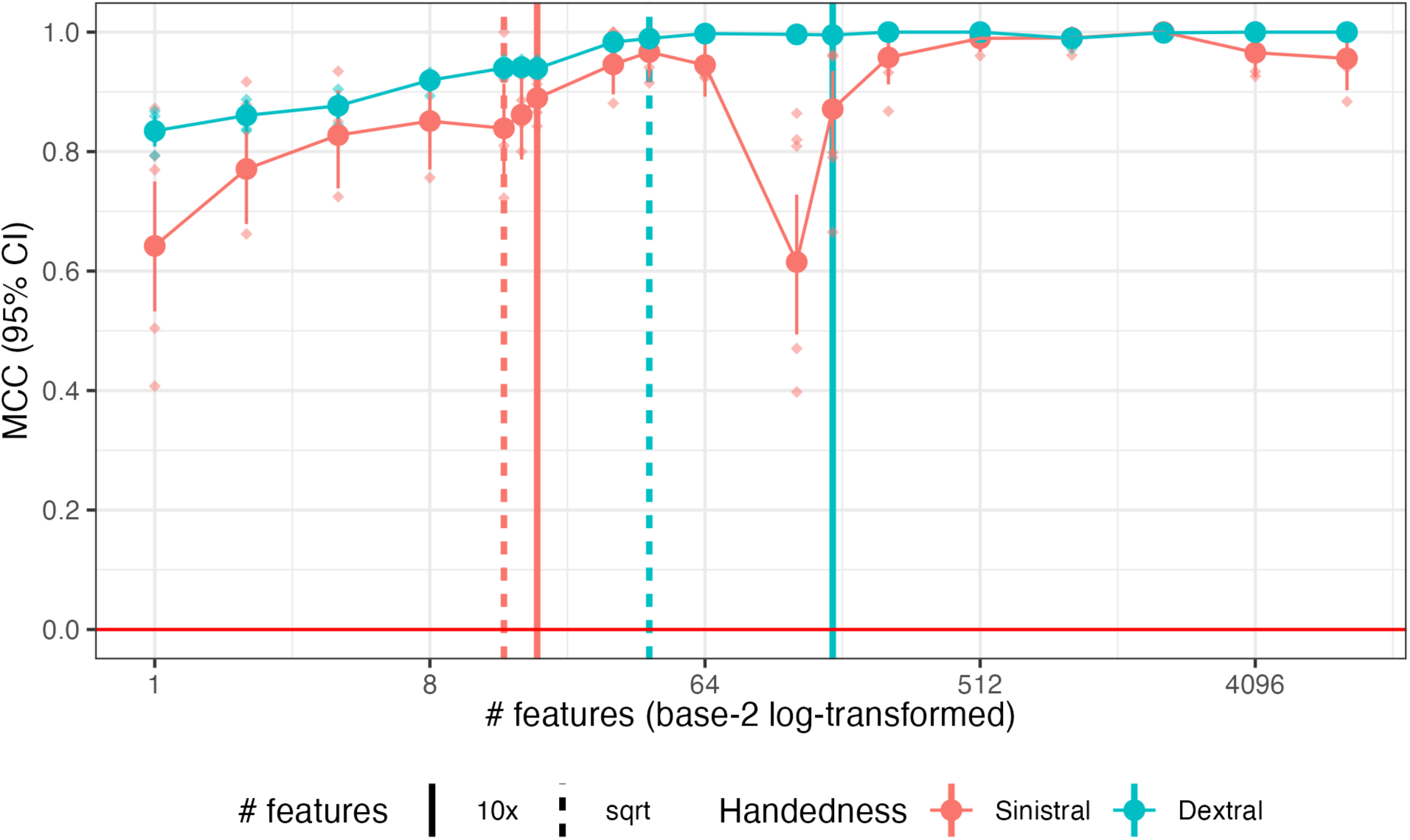
MCC and bootstrapped 95% CI for hemisphere identification by number of features (log-transformed) and by training groups. The five-fold validations are shown as diamonds with the overall MCC as large circles with the confidence interval. Rules of thumb for maximum number of features for each sample are shown in vertical lines.

We tabulate which regions appear in the most important connections for interpretation. The top features based on the *F* classifier as calculated in the dextral and sinistral samples are shown in Table A1.R2. The top 16 features in either classifier are shown, total: 22. Sixteen out of 22 connections involved the perisylvian language area (PSL). No other area appeared more than twice.

**Table A1.R2:** Top features in dextral or sinistral analyses, sorted by their dextral rank.

| Feature | Sinistral rank | Dextral rank | Feature | Sinistral rank | Dextral rank |
| --- | --- | --- | --- | --- | --- |
| 6ma-PSL | 1 | 1 | FOP1-PSL | 3 | 12 |
| IFSa-PSL | 2 | 2 | PGi-PSL | 5 | 13 |
| PF-PSL | 6 | 3 | a47r-45 | 37 | 14 |
| 47l-PSL | 8 | 4 | STSva-PSL | 20 | 15 |
| 47l-45 | 7 | 5 | PoI1-PSL | 13 | 16 |
| 23c-PSL | 21 | 6 | 9m-PSL | 16 | 28 |
| TE1a-PSL | 32 | 7 | PHT-PSL | 10 | 34 |
| MI-PSL | 23 | 8 | PGs-46 | 11 | 36 |
| 7Am-PSL | 12 | 9 | IP1-IFSa | 14 | 46 |
| 8BL-PSL | 39 | 10 | PGs-v23ab | 15 | 47 |
| 46-PSL | 9 | 11 | PGi-8Ad | 4 | 107 |

#### 2.2.3 Parcel symmetrization robustness test

Because the Glasser atlas is not perfectly topologically symmetric, this raises the question of whether the in-built asymmetry in the atlas drives the classification. There is an effect of parcel asymmetry (see Supplemental Material 6), where more asymmetric parcels have higher scores, *B_dextral_ = −0.10, p < .001, B_sinistral_ = −0.12*, *p < .001* and an effect of size, where larger parcels have lower scores, *B_dextral_ = −0.053, p < .001, B_sinistral_ = −0.051, p < .001*, but the effects are small, both *R^2^*s *< 0.015.* While the majority of the top important connections are more asymmetric than average, the vast majority of asymmetric parcels are not highly important, see Supplemental Figure 6.2, suggesting that the classification does rely on lateralized connections, not solely asymmetries introduced by parcel differences.

As a further test, we symmetrize the atlas in both directions and perform the same approach. Hemisphere classification was still perfect in datasets trained and tested on dextrals when using the symmetrized parcels, using both Glasser-LL and Glasser-RR. Classification was slightly lower, but still nearly perfect when trained/tested only on sinistrals, Glasser-LL *MCC = .916 [.857, 975]*; Glasser-RR MCC: *MCC = .916 [.858, 975]*. This provides further evidence that asymmetry in the Glasser parcels do not underlie the classification performance.

We extracted *F* scores from the K-best classifier for each model (Glasser-LL/-RR; dextral/sinistral). The Spearman’s *rho* between features for the models trained on dextrals were *r_LL_ = .55, r_RR_ = .56*, and for the models trained on sinistrals, *r_LL_ = .41, r_RR_ = .43*.

#### 2.2.4 Individual hemisphere scores

Next, we examine how individuals’ hemispheres are placed in linear discriminant space together. Both hemispheres from each participant received a score based on the overall model. LHs predominantly received negative values and RHs, positive values, each centered on ±4.25. We also calculate a “distance” score between each individual’s two hemispheres scores as *distance = RH - LH*, mean = 8.5 (SD: 7.6). These hemisphere scores (“fingerprints”) are inversely correlated with each other, *r_dextral_ = −0.41, p < .001; r_sinistral_ = -.61, p < .001*. In other words, for individuals with LHs that have more negative values, their RHs have more positive scores. In the model trained and tested on all participants, the correlation between individuals’ hemisphere scores was *r_all_ = −0.41, p < .001*.

Furthermore, more strongly right-handed individuals have greater distance between their hemispheres, *β = 0.004, p < .001*, but the effect is slight, *R^2^ = .015.* Because of the common practice that divides handedness into discrete categories like left/right or left/mixed/right, we examine this effect further with segmented linear regression (Muggeo, 2003, 2008) to examine whether distinct groups can be found based on the relation between handedness and hemisphere distance. A one-breakpoint model improves on a straightforward linear regression, *p = .01*, with an improved *R^2^* of .021. However, adding a second breakpoint does not improve over the one-breakpoint model, *p = .94.* In the one-breakpoint model, the breakpoint falls at EHI = 72.7.

These findings show that dextrals have hemispheres that are more distinct from one another compared to sinistrals, evident in both the distance results and in the results that show that their hemisphere scores are less correlated (*|r| = .41*) than sinistrals (*|r| = .61*).

**Figure A1.R3:**
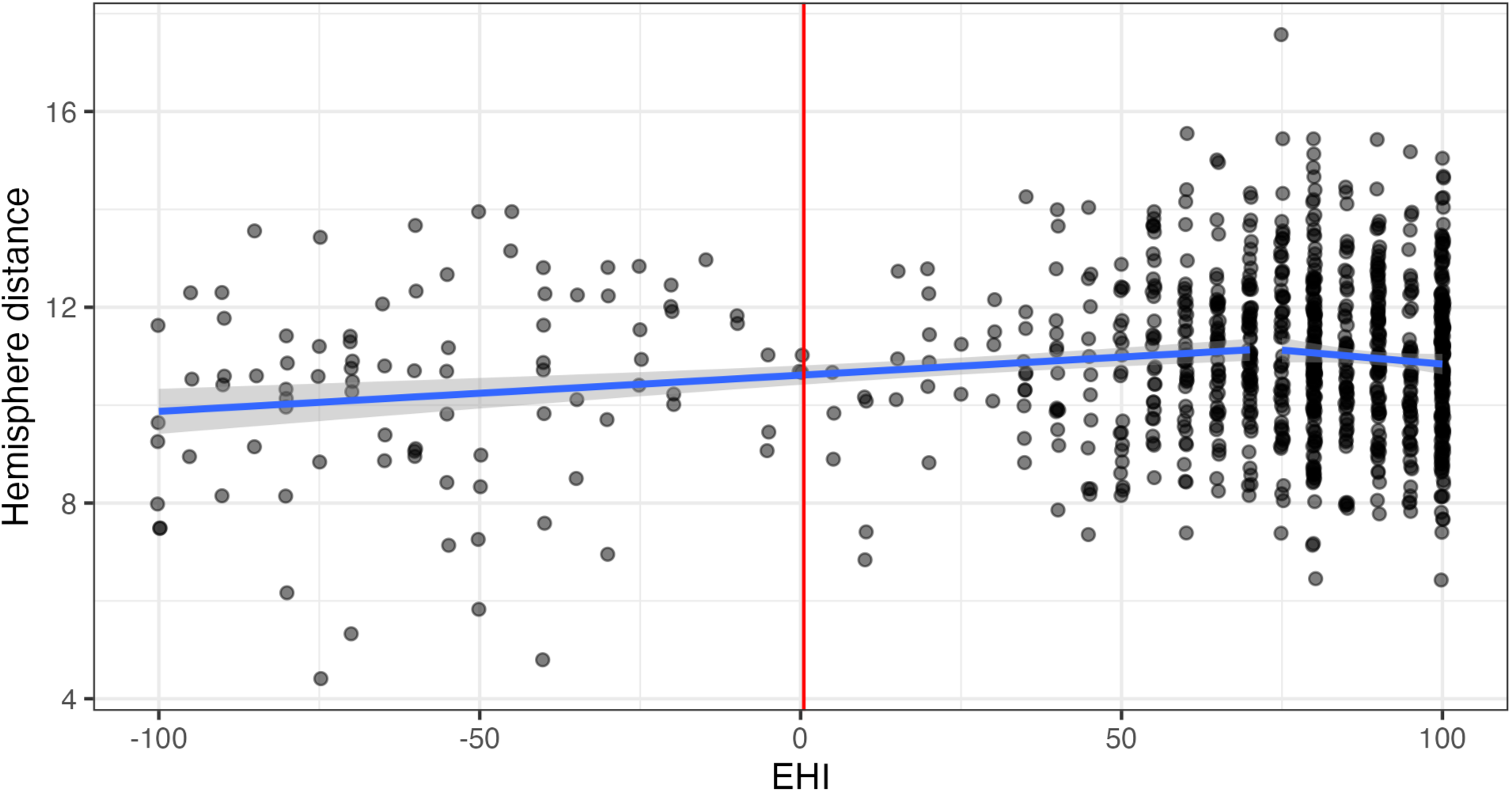
Distance between hemispheres in each dataset. EHI handedness scores are expanded to take up the entire axis for dextrals and sinistrals only for visibility. Note that the magnitude of the hemisphere distance varies.

#### 2.2.5 Feature mapping

We conducted an “enrichment” analysis to identify which connections had a weight greater than expected by chance. Next, we evaluated which networks these connections were between. After bootstrapping for connection significance, 18 connections had simulated *p*-values of 0.0, which were replaced with .0003, as described in the methods. For descriptive purposes, 137 connections had a *p*-value < .01, and 43 had a *p*-values < .001, including the replaced values. See Figure A1.R4, where the three small networks are omitted from A1.R4(B). These results are calculated on the model trained on all available participants (dextrals and sinistrals together).

**Figure A1.R4:**
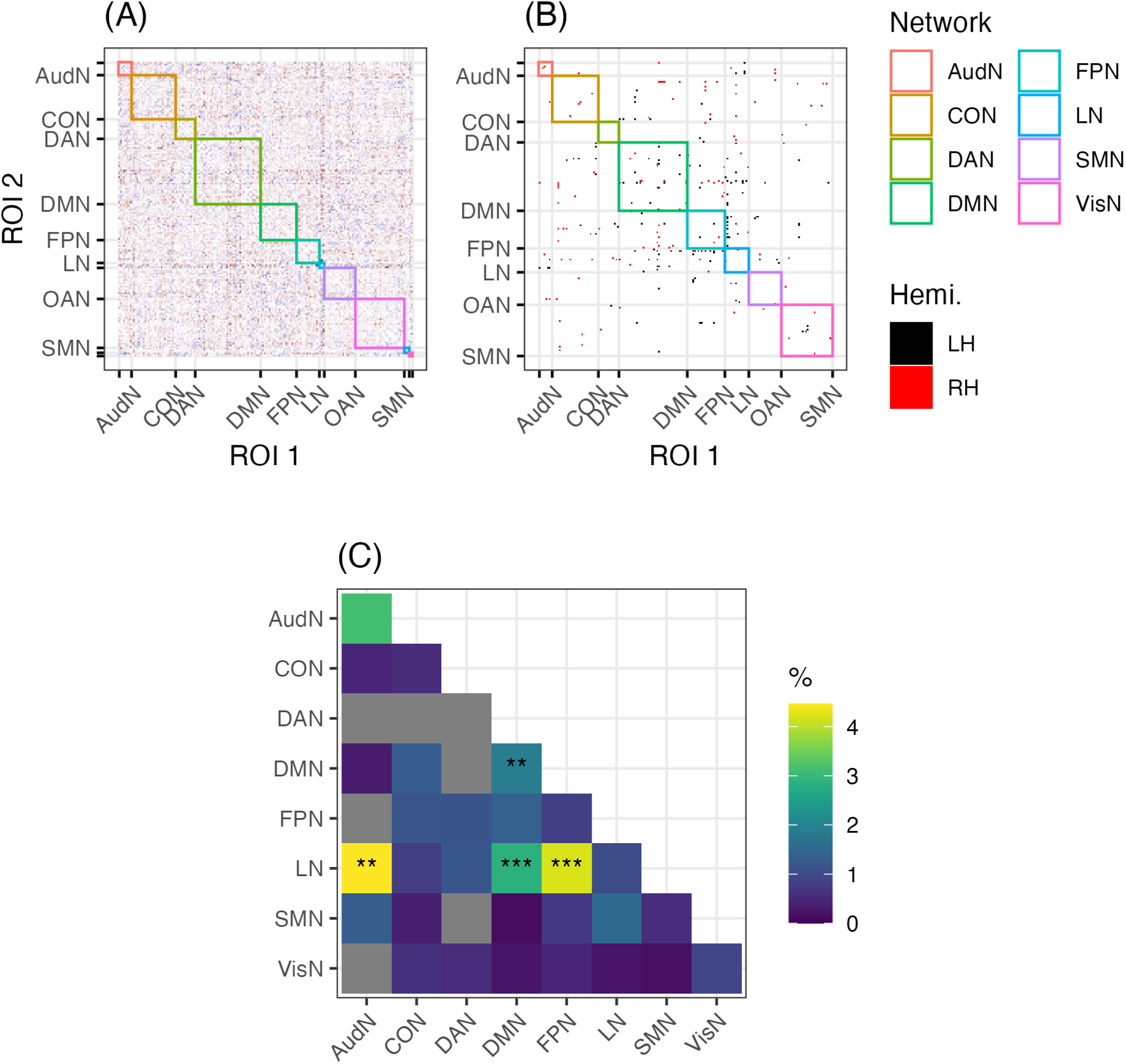
Enrichment results. (A) Scaling values for all connections, by network. (B) Which of these values are higher than expected by chance, colored by which hemisphere has stronger connectivity. (C) Which network pairs (or single networks) have more significant connections than expected by chance: * uncorrected *p* < .05; *** uncorrected *p* < .001. Networks with no significant connections (after step B) are in grey, and networks are colored by the proportion of connections within that group that are significant, even if there aren’t more than expected by chance.

Following FDR correction, the identification of within-DMN connections as important to the model remains significant, *p_FDR_ = .026*, as do the three between-network sets of connections: LN-AudN *p_FDR_ = .026;* LN-DMN *p_FDR_ = .005;* and LN-FPN *p_FDR_ = .005.* Within-LN connections were not important to the model, uncorrected *p = .54*, and neither were within-AudN connections, uncorrected *p = .22*.

For the robustness analyses, see Supplemental Material 7, where the within-DMN, LN-FN, and LN-DMN pattern is robust. The AudN-LN connections do not survive the second robustness analysis, where parcels were removed, but this procedure removed 44% of the already small set of AudN-LN connections (*n_orig_ = 112*), see Supplemental Material 7.2.

### 2.3 Analysis Summary

In this analysis, we show that hemisphere sidedness is trivial to classify, at least in models trained on dextrals. Furthermore, while we primarily test models trained on all 16,110 within-hemisphere connections, models of this sort can be highly successful with only a few hundred selected features. We also show that when an individual’s hemiconnectomes are projected into a one-dimensional “hemisphere axis,” some individuals have hemispheres that are more similar to each other than others do. This difference varies with handedness, with sinistrals having less distinct hemispheres than dextrals, an effect that persists for individuals with EHI scores less than 73.

However, we show that performance is poorer in sinistrals, and that poorer performance is not attributable only to sample size or to the greater variance in absolute handedness score in sinistrals. Secondly, as a check against leakage, when only one hemisphere per participant is included, classification performance is poorer, but this also can be attributable to having reduced the training data by half.

While the connections most important to the classifier are slightly more asymmetric than average, using perfectly mirrored parcellations does not significantly affect classifier performance. This suggests that the model relies on connections that are asymmetric, not only on parcels that happen to be asymmetric between the hemispheres. Indeed, the parcels are asymmetric because the functional and cytoarchitectonic boundaries are thought to be different (Glasser et al., 2016) and using these areas as described likely better captures the actual functional connectivity between regions.

The strongest differences between the hemispheres are found in LN-DMN and LN-FPN connections. The FPN, DMN, and LN are highly related networks. Broadly speaking, the FPN regulates both the DMN and the LN (Gordon et al., 2016). These networks feature less in the lateralization literature (Cole et al., 2015; Hearne et al., 2016; Sheffield et al., 2015), although some authors have reported left-lateralization of the DMN and FPN (Agcaoglu et al., 2015).

The DMN, rather than the LN, was the sole network where the classifier relied on within-network connectivity to predict hemisphere sidedness. This finding recapitulates two previous observations: LN-DMN segregation is stronger in the RH compared to the LH (Mineroff et al., 2018), and the anterior subnetwork of the DMN preferentially connects to the LN, compared to the posterior (Gordon et al., 2020). Greater segregation in the RH is a result of weaker connectivity between networks, and our classifier also relies on LH/RH differences in connectivity between networks, without appealing to graph theory.

## 3 Analysis 2: Handedness classification

### 3.1 Methods

Because of the differences in cortical organization and structural features between dextrals and sinistrals, we sought to ascertain if connectomes could be used to identify handedness as an outcome, in addition to hemisphere sidedness.

The same input and preprocessing steps were taken as described in Analyses 2. However, for determining handedness and not hemisphere polarity, it is now possible to also test the performance of two additional sets of input data: (1) the full connectome (FC), as well as (2) the interhemispheric connections excluded from the hemiconnectomes (the “transconnectome,” or TC). These new featuresets ask whether (1) there is information that distinguishes sinistrals/dextrals in the whole-brain pattern of connectivity; and (2) whether there is information in the transcallosal connections that distinguish sinistrals/dextrals. The FC and TC values were Fisher-Z transformed and mean-centered and normalized across participants.

#### 3.1.1 Class imbalance

Predicting handedness presents an imbalanced-classes problem (Lemaître et al., 2017; Yang & Wu, 2006), because only 10% of the sample is sinistral. Generally speaking, the risk is that the model predicts all observations as belonging to the majority class (i.e., dextrals), which could reach accuracies of 90%, while completely failing one group (sinistrals). MCC ameliorates this risk, by assigning an MCC of 0 if any class, no matter the size, is completely incorrect (see Supplemental Material 4).

One method for addressing imbalance data is oversampling, i.e. duplicating rows until equal class sizes are achieved. We created a dataset (Hemiconnectome-Oversampled; HCO) that oversampled sinistral participants using the upsample() function, from the R package groupdata2 (ver. 2.0.5; Olsen, 2024). These values were scaled as in Analysis 2 and then oversampled.

#### 3.1.2 Outcomes

For the hemiconnectomes (HC, LgHC, and HCO), we predict a four-way outcome: LH vs. RH crossed with sinistral vs. dextral. We then extract handedness and sidedness prediction from the four-way prediction and examine performance for the two sub-predictions alongside overall prediction. For FC and TC, we predict only handedness because these data include both hemispheres, and neither hemisphere, respectively. In this situation, MCC is also useful to compare between the four-class and two-class outcomes, whereas using accuracy, the random prediction value would be different: .25 and .5, respectively. Lastly, handedness is slightly related to sex, and as a robustness test, we add sex to the outcome (creating an eight-way classification problem).

### 3.2 Results

#### 3.2.1 Handedness identification

As in Analysis 1, we start by reporting the model success rates. None of the LDA models (*n = 75*) succeeded in identifying handedness, with the overall MCC falling between [.058, .285], with none of the handedness-prediction models passing the “good” threshold of MCC > .4, see Figure A2.R1 (note the change in color scheme). However, hemisphere identifications for the models (where it is possible) remained > .99.

**Figure A2.R1:**
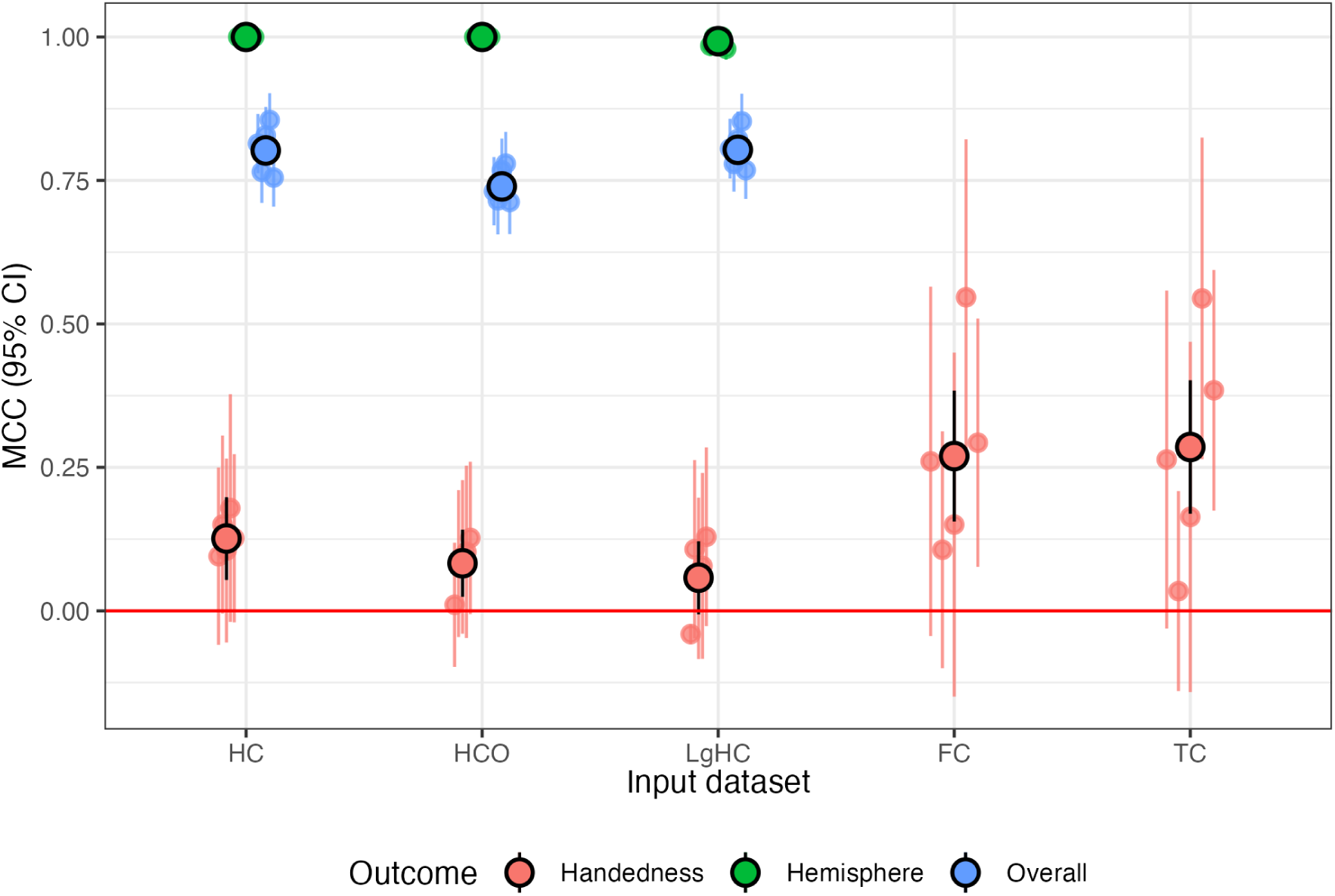
MCCs for the handedness prediction across folds and input data. The large, outlined points display the MCC calculated on the overall sample, and the smaller points, the MCC on a per-fold basis. HC: hemiconnectomes, HCO: hemiconnectomes oversampled; LgHC: language task hemiconnectomes; FC: full connectomes; TC: transconnectomes.

As before, we perform an ANOVA to examine which input dataset choices affect the classification. In a two-way ANOVA across the handedness prediction in all five input datasets, there was an effect of fold, *F(4) = 3.3, p = .037*, and of input dataset, *F(4) = 5.0, p = .008.* None of the pairwise comparisons for folds were significant after adjustment. Out of the 10 pairwise comparisons for the input datasets, the language connectome (LgHC) underperformed the two non-hemisphere models, i.e., LgHC < FC, *Δ = -.21, p = .04*, and LgHC < TC, *Δ = -.21, p = .03*.

However, none of these models can be described as accurately classifying handedness; only two folds reached even the previously described MCC > .4 “good” threshold, which did not generalize to the rest of the FC or TC folds. Trends in the NN and SVC models were similar, handedness MCC range for NN: [.077, .188], for SVC: [.088, .274], all hemisphere MCCs > .981, see Supplemental Figure 5.1.

#### 3.2.2 Gender control

Crossing gender with the outcome variables did not improve the handedness classification accuracy. In a three-way ANOVA controlling for fold and input dataset (FC, TC, HC), there was no effect of including gender, *F(1) = 2.32, p = .14*. However, classification of gender itself was fairly high, MCC range: [.770, .873], which is consistent with prior work (Weber et al., 2022; C. Zhang et al., 2018). See also Supplemental Material 8.

#### 3.2.3 Feature exploration

As with the hemisphere models, we explored the LDA feature values to understand how handedness projects onto a lower-dimensional space. Because this model has four outcomes, there were three LDs, compared to one LD (based on two classes: LH/RH) previously. First, we plot each individual’s first (LD1) and second linear discriminants (LD2) against each other. These LDs explained 95% and 3% of the variance, respectively. Sinistrals and dextrals cannot be separated on these axes, see Figure A2.R2.

Sinistrals have hemispheres that are, on average, closer in LD1 (or “hemisphere space”), i.e., the horizontal distance between the rectangular centroid markers in Figure A2.R2 is significant in both hemispheres, tested separately with a *t*-test: LH: *t(104.8) = 3.7, d = −0.54, p < .001*; RH: *t(106.9) = −4.4, d = .47, p < .001*, but there is no difference in LD2 scores, both *p* values > .2. This replicates the results of Analysis 1, where sinistrals had more similar hemispheres, and shows this effect is unaffected by the cross with handedness.

**Figure A2.R2:**
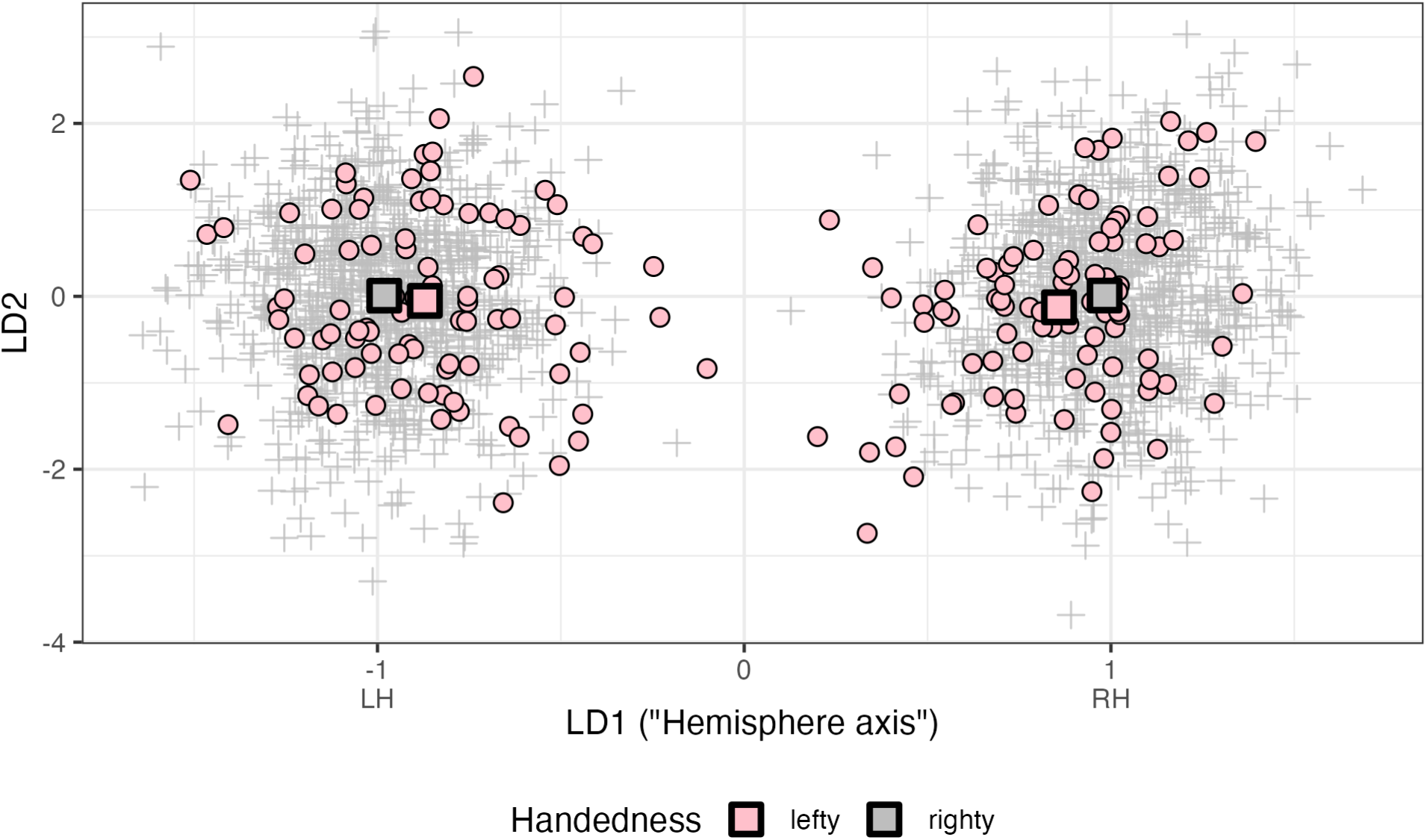
First and second linear discriminants for the four-outcome model predicting hemisphere and handedness status.

Because including handedness did not affect the hemisphere classification success rate, we did not repeat the LD1-against-EHI analyses shown in section 2.2.4, nor the investigation of network enrichment, as there appears to be no change. Because the handedness classification did not succeed, there are no meaningful features to explore.

### 3.3 Analysis Summary

Overall, our functional connectivity analyses fail to distinguish sinistrals from dextrals. This failure cannot be attributed solely to the smaller training group, as hemisphere classification on dextrals succeeds with groups of similar size. Although the literature has found structural differences between sinistrals and dextrals (Caeyenberghs & Leemans, 2014; Sha, Schijven, et al., 2021), our findings suggest that these differences are not reflected in functional connectivity.

Furthermore, because men are 22% more likely to be sinistral (men: 11.6%) than women (9.5%), we attempted to control for participant sex in the classification; but this did not change the classifier success rate.

That said, the models were still able to identify hemisphere sidedness with near-perfect accuracy, even in the sinistral participants (and even when the handedness was identified incorrectly), suggesting that the differences between the hemispheres are larger and more stable than those between dextrals and sinistrals.

## 4 Discussion

Our goals were to use supervised learning models to test whether it was possible to identify a hemisphere as left or right, or belonging to a sinistral or dextral person based only on its within-hemisphere connectivity. All three types of models (LDA, NN, SVC) almost perfectly identify the hemispheres in dextrals across multiple types of functional connectivity data from different task states, with slightly lower, but still high rates in sinistrals as well. To our knowledge, this is the first demonstration that hemisphere sidedness can be identified from resting-state functional connectivity.

### 4.1 Hemisphere identification

In linear-discriminant “hemisphere space,” we observe that, on average, sinistrals have hemispheres that are more similar to one another than dextrals do, although none are so similar that the LDA classifiers misclassify it. Secondly, generally speaking, using data collected during a language task (but treated as resting-state) did not improve performance, although this may be attributable to the reduced acquisition time, and not a lack of task-induced connectivity changes. It is possible that connectivity changes induced in a longer task may prove helpful in distinguishing sinistrals and dextrals.

Our success in identification is supported by the literature identifying differences in structure (Sha, Schijven, et al., 2021) and structural connectivity between the hemispheres (Caeyenberghs & Leemans, 2014; D. Li et al., 2019; M. Li et al., 2014), including in non-human primates (Iturria-Medina et al., 2011).

However, the difference between the hemispheres is not isolated to the language network. Rather, the strongest differences between the hemispheres are found in LN-DMN and LN-FPN connections. The FPN, DMN, and LN are highly related networks. Broadly speaking, the FPN regulates both the DMN and the LN (Gordon et al., 2020), and connectivity between these networks and between these networks and the rest of the brain is positively correlated with measures of intelligence (Marek & Dosenbach, 2018). Based on this work, an investigation of hemispheric difference and intelligence could serve as a test of external validity.

The DMN, rather than the LN, was the sole network where the classifier relied on within-network connectivity to predict sidedness, in dextrals, sinistrals and the dextral+sinistral analyses. The DMN and LN are strongly segregated from one another in task activations, an effect which is even stronger in RH compared to the LH (Mineroff et al., 2018). Although we do not investigate subnetworks of the large-scale RSNs in this analysis, the anterior subnetwork of the DMN preferentially connects to the LN, compared to the posterior (Gordon et al., 2020). This asymmetry in connectivity is reflected in broadly more efficient and less random organization in the LH compared to the RH (Caeyenberghs & Leemans, 2014; Dennis et al., 2013). These differences in efficiency are present from birth (Ratnarajah et al., 2013), which suggests genetic biases for hemispheric organization. Network-based statistics like segregation and efficiency are based on edge weights, i.e., parcel-to-parcel correlations, so it is unsurprising to recapitulate these features here.

### 4.2 Handedness identification

The distance between hemispheres scales with a continuous measure of handedness, with more sinistral participants having more similar hemispheres, for individuals with an EHI value < 73. This value suggests a neural basis for the cutoff between sinistral (or nondextral) and dextral brains, which is higher than usually suggested (Papadatou-Pastou et al., 2020).

We did not find neural support for three handedness groups. In contrast, the psychometric literature has often found bases for dividing handedness into three groups (Dragovic, 2004; Papadatou-Pastou et al., 2020; Peters, 1992), but some authors find two groups (Büsch et al., 2010). For a two-group division (dextral vs. adextral), an EHI cutoff between +30 and +40 is common, see Przybylski and Kroliczak (2023) and citations within. Our value (+73) is much greater than this. But the lack of a second discontinuity between “sinistrals” and “mixed-handedness” may be affected by the rate of adextral participants in our sample.

That said, unlike hemispheres, which have no intermediate kinds, separating individuals with similar EHIs of, e.g., −5 and +5 into different groups — as was done here — may impair model performance. We did not test “strong handedness” groups, i.e. only sinistrals < −30 vs. dextrals > +30, in order to not decrease the size of the already-small (*n = 74*) sinistral training groups.

One possible outcome was that strong dextrals and strong sinistrals (i.e., high absolute EHI values) patterned together, with differences between these groups and ambidextrous/mixed participants. We did not find such a U-shaped curve. This lack of curve implies that greater separation between the hemispheres is associated with right-handedness, and not associated with a strong hand preference in general. That said, as with most resting-state studies (Marek et al., 2022), the effects are very small, explaining only a few percent of the variance.

Finally, from the hemisphere-by-handedness LDA, the second and third linear discriminants provide hints that sinistrals are organized on one extreme of the norm, even if the pattern wasn’t sufficient to classify a connectome. It may be the case, therefore, that an analysis based on different methods of characterizing brain activity may be more fruitful in the classification problem. Using derived values, like network-based statistics (c.f. Caeyenberghs & Leemans, 2014) or differences between matched features may permit handedness classification.

### 4.3 Analytic decisions

We used a single parcellation scheme that had some advantages, namely that there were the same (albeit, non-homotopic) parcels in each hemisphere. Other schemes have found different parcels (in number and extent) in the two hemispheres, for example, Schaefer and colleagues (2018) parcellated the cortex into 10 levels of granularity between 50 and 500 parcels in each hemisphere, although not with one-to-one transcallosal correspondence. It may be possible that a different scheme might better capture variability, especially in the RH, that is attributable to handedness.

Likewise, using a single parcellation scheme for a large number of participants is a common analysis technique, but sinistrals are typified by their heterogeneity. That is, sinistrals are less consistently lateralized for language (Knecht et al., 2000; Willems et al., 2014) and motor control (Willems et al., 2014). If this broad-scale heterogeneity is reflected in parcel-level heterogeneity, a parcel applied to a sinistral brain may, on average, contain signals from more nearby, but unrelated regions than the same parcel applied to a dextral brain. As a consequence, the signal-to-noise ratio in a given parcel may be lower in sinistrals, harming model performance.

### 4.6 Future research in hemisphere distinctiveness

The creation of a scalar to measure the distance between hemispheres is advantageous for exploration of experiential effects on hemisphere organization. We propose three main avenues for future work in hemispheric distinctiveness. These areas are: hemispheric reorganization following stroke, typical development, and lifelong experience with a dominant hand.

It also remains an open question how the uninjured hemisphere reorganizes after a unilateral cortical stroke. DMN and FPN connectivity can be disrupted following a stroke, even if the lesion is outside of the network (Dacosta-Aguayo et al., 2015; Tuladhar et al., 2013), with changes in disruption patterns from the acute to chronic phase (Z. Zhang, 2025). Conversely, damage to LH language areas typically causes aphasia, and damage to other regions, including the RH LN, does not. In people with aphasia, the RH becomes modestly more involved in language processing, through increased recruitment of RH language regions, and recruitment of new RH areas, depending on lesion characteristics (Turkeltaub et al., 2025; Wilson & Schneck, 2021). Connections between the damaged LN and the FPN improve outcomes (Billot & Kiran, 2024).

Our work shows that different patterns of LN-FPN and LN-DMN connections are among the most distinctive elements of the LH. As such, we can ask whether intact RHs tend to become more like LHs in the chronic period. While connectivity work has evaluated the connections between these networks, our protocol provides a simple, easily interpretable number (or set of numbers) that could be used to quantify “how much” an RH looks like a LH. Thus, we plan to apply this analysis to stroke connectomes to quantify gestalt hemispheric reorganization longitudinally over recovery. The hemispheric fingerprinting approach would also be appropriate to examine hemispheric reorganization in other disease processes where unilateral dysfunction has been shown to result in contralateral compensation, such as brain tumors and focal epilepsies.

Secondly, we also propose to use this model longitudinally. Is the “distance” between hemispheres innate at birth, i.e., differing by handedness groups, or does it emerge over time? Evidence from language laterality suggests that language is more bilateral in childhood (Olulade et al., 2020; Szaflarski, Schmithorst, et al., 2006), thus we would expect the difference between the hemispheres to emerge over time, but the exact timing remains an empirical question.

However, a developmental study could show the direction of the effect: Do more weakly lateralized brains at birth become left-handed, and strongly lateralized brains become right-handed? Or does a third factor determine whether weakly lateralized brains at birth (1) become distinct-and-dextral or (2) exhibit less change and become sinistral? This study would test whether there is a causal effect of handedness on brain organization.

While a decrease in bilaterality in development seems to occur, the reverse effect has been proposed in aging. This model is known as “hemispheric asymmetry reduction in older adults” (HAROLD; Cabeza et al., 2002), where functional lateralization decreases with time. While some studies have failed to support this hypothesis (Chang et al., 2026; Meunier et al., 2014; Nenert et al., 2017), our model could provide a simple test of this hypothesis in similar open datasets.

Finally, writing is especially interesting as a motor behavior because there is a multicultural history of teaching natural sinistrals to write both alphabetic and logographic scripts with their non-dominant (i.e., right) hand (Guber, 2019; Kushner, 2012, 2013; Papadatou-Pastou et al., 2020). These “converted” sinistrals are aging, as the practice has largely fallen out of favor, ending in the U.S. in the 1970s (Hugdahl et al., 1993). However, there is structural (Klöppel et al., 2007, 2010) and motor-task evidence (Grabowska et al., 2012; Siebner et al., 2002) that this population differs cortically and subcortically from dextrals and non-converted sinistrals, primarily in the direction of increased bilaterality.

With respect to hand use, after dominant right-hand amputation, amputees show changes in control pathways and increase in LH activation for control of the non-amputated left hand (Philip & Frey, 2014). Applying this algorithm to these populations would also provide a natural experiment testing the effects of changes in motor dominance on cortex-wide organization; for example, are these individuals’ cortical organization much more like a strong dextral than their handedness would otherwise predict?

## 5 Conclusions

We find that it is trivial to identify the sidedness of a hemisphere given its functional connectivity patterns during rest. This classification relies on connections between the language network and the default mode and frontoparietal networks, but not connections within the language network itself. In contrast, we could not reliably classify a hemisphere as belonging to a sinistral or dextral.

However, analyses of the difference *between* hemispheres suggests that there is an effect of handedness on hemispheric similarity, for all but the strongest right handers. With these findings, we can ask how the separation between the hemispheres emerges over development, does the distance increase with age, or is it a stable trait? The approach to quantifying the similarity between hemispheres we have described here opens up new avenues of research to investigate the nature of brain lateralization and situations in which it may differ from the strong lateralization observed in typical dextral adults, including recovery after cortical injury, development, and changes in fine motor use.

## 6 Ending Sections

### 6.1 Data and code availability

All data are publicly available through the Human Connectome Project or Zenodo (for the parcel-to-network assignments). Analysis code is available at https://github.com/TrevorKMDay/hemidentification

### 6.2 Author contributions

● TKMD: Conceptualization, data management, formal analysis, investigation, methodology, project administration, software, visualization, writing - original draft, writing - review & editing;
● PET: Conceptualization, funding acquisition, resources, supervision, writing - review & editing;
● ELN: Conceptualization, funding acquisition, resources, supervision, writing - review & editing;
● ATD: Formal analysis, methodology, project administration, supervision, writing - review & editing.

### 6.3 Funding

This work was supported by NIH T32DC019481 and the MedStar National Rehabilitation Network (TKMD), NIH R01 DC014960 (PET), NSF BCS-2318609 and the Bergeron Endowed Professorship (ELN), and NIH R00 DC018828 (ATD).

## 6.5 Acknowledgements

The authors wish to thank Andrew Hannum, PhD, and Saúl Blanco, PhD, for their input during the planning of these analyses and Kyle Shattuck, PhD, for his computational assistance.

## Footnotes

1 Regions that are part of the LH LN, but a different network in the RH, by RH network: CON (region PSL), pMN (STV), FPN (44, IFSp), and DMN (STSda).

2 https://infer.tidymodels.org/reference/get_p_value.html

## References

Agcaoglu, O., Miller, R., Damaraju, E., Rashid, B., Bustillo, J., Cetin, M. S., Van Erp, T. G. M., McEwen, S., Preda, A., Ford, J. M., Lim, K. O., Manoach, D. S., Mathalon, D. H., Potkin, S. G., & Calhoun, V. D. (2018). Decreased hemispheric connectivity and decreased intra- and inter-hemisphere asymmetry of resting state functional network connectivity in schizophrenia. Brain Imaging and Behavior, 12(3), 615–630. 10.1007/s11682-017-9718-7

Agcaoglu, O., Miller, R., Mayer, A. R., Hugdahl, K., & Calhoun, V. D. (2015). Lateralization of resting state networks and relationship to age and gender. NeuroImage, 104, 310–325. 10.1016/j.neuroimage.2014.09.001

Barch, D. M., Burgess, G. C., Harms, M. P., Petersen, S. E., Schlaggar, B. L., Corbetta, M., Glasser, M. F., Curtiss, S., Dixit, S., Feldt, C., Nolan, D., Bryant, E., Hartley, T., Footer, O., Bjork, J. M., Poldrack, R., Smith, S., Johansen-Berg, H., Snyder, A. Z., & Van Essen, D. C. (2013). Function in the human connectome: Task-fMRI and individual differences in behavior. NeuroImage, Mapping the Connectome, 80, 169–189. 10.1016/j.neuroimage.2013.05.033

Bernard, F., Lemee, J.-M., Mazerand, E., Leiber, L.-M., Menei, P., & Ter Minassian, A. (2020). The ventral attention network: The mirror of the language network in the right brain hemisphere. Journal of Anatomy, 237(4), 632–642. 10.1111/joa.13223

Billot, A., & Kiran, S. (2024). Disentangling neuroplasticity mechanisms in post-stroke language recovery. Brain and Language, 251, 105381. 10.1016/j.bandl.2024.105381

Binder, J. R., Gross, W. L., Allendorfer, J. B., Bonilha, L., Chapin, J., Edwards, J. C., Grabowski, T. J., Langfitt, J. T., Loring, D. W., Lowe, M. J., Koenig, K., Morgan, P. S., Ojemann, J. G., Rorden, C., Szaflarski, J. P., Tivarus, M. E., & Weaver, K. E. (2011). Mapping anterior temporal lobe language areas with fMRI: A multicenter normative study. NeuroImage, 54(2), 1465–1475. 10.1016/j.neuroimage.2010.09.048

Bishop, D. V. M. (2013). Cerebral Asymmetry and Language Development: Cause, Correlate, or Consequence? Science, 340(6138), 1230531. 10.1126/science.1230531

Büsch, D., Hagemann, N., & Bender, N. (2010). The dimensionality of the Edinburgh Handedness Inventory: An analysis with models of the item response theory. Laterality, 15(6), 610–628. 10.1080/13576500903081806

Caeyenberghs, K., & Leemans, A. (2014). Hemispheric lateralization of topological organization in structural brain networks. Human Brain Mapping, 35(9), 4944–4957. 10.1002/hbm.22524

Chang, E. H., Seydell-Greenwald, A., Day, T. K. M., Nisbet, D., & Turkeltaub, P. E. (2026). Language remains strongly left-lateralized in older adults: A cross-sectional and longitudinal study (p. 2026.07.17.739182). bioRxiv. 10.64898/2026.07.17.739182

Chicco, D., & Jurman, G. (2023). The Matthews correlation coefficient (MCC) should replace the ROC AUC as the standard metric for assessing binary classification. BioData Mining, 16(1), 4. 10.1186/s13040-023-00322-4

Ciric, R., Rosen, A. F. G., Erus, G., Cieslak, M., Adebimpe, A., Cook, P. A., Bassett, D. S., Davatzikos, C., Wolf, D. H., & Satterthwaite, T. D. (2018). Mitigating head motion artifact in functional connectivity MRI. Nature Protocols, 13(12), 2801–2826. 10.1038/s41596-018-0065-y

Cole, M. W., & Ito, T. (2018). *ColeLab/ColeAnticevicNetPartition: First public release of The Cole-Anticevic Brain-wide Network Partition (CAB-NP)* (Version v1.0.5-major) [Computer software]. Zenodo. 10.5281/zenodo.1455791

Cole, M. W., Ito, T., & Braver, T. S. (2015). Lateral Prefrontal Cortex Contributes to Fluid Intelligence Through Multinetwork Connectivity. Brain Connectivity, 5(8), 497–504. 10.1089/brain.2015.0357

Corbetta, M., Patel, G., & Shulman, G. L. (2008). The Reorienting System of the Human Brain: From Environment to Theory of Mind. Neuron, 58(3), 306–324. 10.1016/j.neuron.2008.04.017

Corbetta, M., & Shulman, G. L. (2002). Control of goal-directed and stimulus-driven attention in the brain. Nature Reviews. Neuroscience, 3(3), 201–215. 10.1038/nrn755

Corbetta, M., & Shulman, G. L. (2011). Spatial Neglect and Attention Networks. Annual Review of Neuroscience, 34(1), 569–599. 10.1146/annurev-neuro-061010-113731

Couch, S., Bray, A., Ismay, C., Chasnovski, E., Baumer, B., & Çetinkaya-Rundel, M. (2021). infer: An R package for tidyverse-friendly statistical inference. Journal of Open Source Software, 6(65), 3661. 10.21105/joss.03661

Crow, T. J. (2000). Schizophrenia as the price that Homo sapiens pays for language: A resolution of the central paradox in the origin of the species. Brain Research Reviews, 31(2), 118–129. 10.1016/S0165-0173(99)00029-6

Dacosta-Aguayo, R., Graña, M., Iturria-Medina, Y., Fernández-Andújar, M., López-Cancio, E., Cáceres, C., Bargalló, N., Barrios, M., Clemente, I., Toran, P., Forés, R., Dávalos, A., Auer, T., & Mataró, M. (2015). Impairment of functional integration of the default mode network correlates with cognitive outcome at three months after stroke. Human Brain Mapping, 36(2), 577–590. 10.1002/hbm.22648

Day, T. K. M. (2024). Resting-state and task functional magnetic resonance imaging correlates of early language development in temporal and frontal lobe [Doctoral dissertation]. University of Minnesota.

Day, T. K. M. (2026). glasser_LR_overlap [Shell]. https://github.com/TrevorKMDay/glasser_LR_overlap (Original work published 2026)

Dennis, E. L., Jahanshad, N., McMahon, K. L., de Zubicaray, G. I., Martin, N. G., Hickie, I. B., Toga, A. W., Wright, M. J., & Thompson, P. M. (2013). Development of brain structural connectivity between ages 12 and 30: A 4-Tesla diffusion imaging study in 439 adolescents and adults. NeuroImage, 64, 671–684. 10.1016/j.neuroimage.2012.09.004

Dewarrat, G. M., Annoni, J.-M., Fornari, E., Carota, A., Bogousslavsky, J., & Maeder, P. (2009). Acute aphasia after right hemisphere stroke. Journal of Neurology, 256(9), 1461–1467. 10.1007/s00415-009-5137-z

Dragovic, M. (2004). Categorization and validation of handedness using latent class analysis. Acta Neuropsychiatrica, 16(4), 212–218. 10.1111/j.0924-2708.2004.00087.x

Eggebrecht, A. T., Elison, J. T., Feczko, E., Todorov, A., Wolff, J. J., Kandala, S., Adams, C. M., Snyder, A. Z., Lewis, J. D., Estes, A. M., Zwaigenbaum, L., Botteron, K. N., McKinstry, R. C., Constantino, J. N., Evans, A., Hazlett, H. C., Dager, S., Paterson, S. J., Schultz, R. T.,…Pruett, J. R. (2017). Joint Attention and Brain Functional Connectivity in Infants and Toddlers. Cerebral Cortex. 10.1093/cercor/bhw403

Finn, E. S., Shen, X., Scheinost, D., Rosenberg, M. D., Huang, J., Chun, M. M., Papademetris, X., & Constable, R. T. (2015). Functional connectome fingerprinting: Identifying individuals using patterns of brain connectivity. Nature Neuroscience, 18(11), 1664–1671. 10.1038/nn.4135

Floris, D. L., Wolfers, T., Zabihi, M., Holz, N. E., Zwiers, M. P., Charman, T., Tillmann, J., Ecker, C., Dell’Acqua, F., Banaschewski, T., Moessnang, C., Baron-Cohen, S., Holt, R., Durston, S., Loth, E., Murphy, D. G. M., Marquand, A., Buitelaar, J. K., Beckmann, C. F.,…Zwiers, M. P. (2020). Atypical Brain Asymmetry in Autism—A Candidate for Clinically Meaningful Stratification. Biological Psychiatry: Cognitive Neuroscience and Neuroimaging, S2451902220302433. 10.1016/j.bpsc.2020.08.008

Glasser, M. F., Coalson, T. S., Robinson, E. C., Hacker, C. D., Harwell, J., Yacoub, E., Ugurbil, K., Andersson, J., Beckmann, C. F., Jenkinson, M., Smith, S. M., & Van Essen, D. C. (2016). A multi-modal parcellation of human cerebral cortex. Nature, 536(7615), Article 7615. 10.1038/nature18933

Glasser, M. F., & Rilling, J. K. (2008). DTI Tractography of the Human Brain’s Language Pathways. Cerebral Cortex, 18(11), 2471–2482. 10.1093/cercor/bhn011

Glasser, M. F., Sotiropoulos, S. N., Wilson, J. A., Coalson, T. S., Fischl, B., Andersson, J. L., Xu, J., Jbabdi, S., Webster, M., Polimeni, J. R., Van Essen, D. C., Jenkinson, M., & WU-Minn HCP Consortium. (2013). The minimal preprocessing pipelines for the Human Connectome Project. NeuroImage, 80, 105–124. 10.1016/j.neuroimage.2013.04.127

Gordon, E. M., Laumann, T. O., Adeyemo, B., Huckins, J. F., Kelley, W. M., & Petersen, S. E. (2016). Generation and Evaluation of a Cortical Area Parcellation from Resting-State Correlations. Cerebral Cortex, 26(1), 288–303. 10.1093/cercor/bhu239

Gordon, E. M., Laumann, T. O., Marek, S., Raut, R. V., Gratton, C., Newbold, D. J., Greene, D. J., Coalson, R. S., Snyder, A. Z., Schlaggar, B. L., Petersen, S. E., Dosenbach, N. U. F., & Nelson, S. M. (2020). Default-mode network streams for coupling to language and control systems. Proceedings of the National Academy of Sciences, 117(29), 17308–17319. 10.1073/pnas.2005238117

Grabowska, A., Gut, M., Binder, M., Forsberg, L., Rymarczyk, K., & Urbanik, A. (2012). Switching handedness: fMRI study of hand motor control in right-handers, left-handers and converted left-handers. Acta Neurobiologiae Experimentalis, 72(4), 439–451. 10.55782/ane-2012-1914

Guber, R. (2019). Making it right? Social norms, handwriting and human capital. Labour Economics, 56, 44–57. 10.1016/j.labeco.2018.11.005

Hale, T. S., Bookheimer, S., McGough, J. J., Phillips, J. M., & McCracken, J. T. (2007). Atypical Brain Activation During Simple & Complex Levels of Processing in Adult ADHD: An fMRI Study. Journal of Attention Disorders, 11(2), 125–139. 10.1177/1087054706294101

Hale, T. S., McCracken, J. T., McGough, J. J., Smalley, S. L., Phillips, J. M., & Zaidel, E. (2005). Impaired linguistic processing and atypical brain laterality in adults with ADHD. Clinical Neuroscience Research, Attention Deficit Hyperactivity Disorder (ADHD), 5(5), 255–263. 10.1016/j.cnr.2005.09.006

Hannum, A., Lopez, M. A., Blanco, S. A., & Betzel, R. F. (2023). High-accuracy machine learning techniques for functional connectome fingerprinting and cognitive state decoding. Human Brain Mapping, 44(16), 5294–5308. 10.1002/hbm.26423

Hearne, L. J., Mattingley, J. B., & Cocchi, L. (2016). Functional brain networks related to individual differences in human intelligence at rest. Scientific Reports, 6(1), 32328. 10.1038/srep32328

Hermosillo, R. J., Moore, L. A., Feczko, E., Miranda-Domínguez, Ó., Pines, A., Dworetsky, A., Conan, G., Mooney, M. A., Randolph, A., & Graham, A. (2024). A precision functional atlas of personalized network topography and probabilities. Nature Neuroscience, 1–14.

Hesterberg, T. (2014). *What Teachers Should Know about the Bootstrap: Resampling in the Undergraduate Statistics Curriculum* (arXiv:1411.5279). arXiv. 10.48550/arXiv.1411.5279

Hua, J., Xiong, Z., Lowey, J., Suh, E., & Dougherty, E. R. (2005). Optimal number of features as a function of sample size for various classification rules. Bioinformatics, 21(8), 1509–1515. 10.1093/bioinformatics/bti171

Hugdahl, K., Satz, P., Mitrushina, M., & Miller, E. N. (1993). Left-handedness and old age: Do left-handers die earlier? Neuropsychologia, 31(4), 325–333. 10.1016/0028-3932(93)90156-T

Iturria-Medina, Y., Pérez Fernández, A., Morris, D. M., Canales-Rodríguez, E. J., Haroon, H. A., García Pentón, L., Augath, M., Galán García, L., Logothetis, N., Parker, G. J. M., & Melie-García, L. (2011). Brain Hemispheric Structural Efficiency and Interconnectivity Rightward Asymmetry in Human and Nonhuman Primates. Cerebral Cortex, 21(1), 56–67. 10.1093/cercor/bhq058

Ji, J. L., Spronk, M., Kulkarni, K., Repovš, G., Anticevic, A., & Cole, M. W. (2019). Mapping the human brain’s cortical-subcortical functional network organization. NeuroImage, 185, 35–57. 10.1016/j.neuroimage.2018.10.006

Kanwisher, N., & Yovel, G. (2006). The fusiform face area: A cortical region specialized for the perception of faces. Philosophical Transactions of the Royal Society B: Biological Sciences, 361(1476), 2109–2128. 10.1098/rstb.2006.1934

Kardan, O., Kaplan, S., Wheelock, M. D., Feczko, E., Day, T. K. M., Miranda-Domínguez, Ó., Meyer, D., Eggebrecht, A. T., Moore, L. A., Sung, S., Chamberlain, T. A., Earl, E., Snider, K., Graham, A., Berman, M. G., Uğurbil, K., Yacoub, E., Elison, J. T., Smyser, C. D.,…Rosenberg, M. D. (2022). Resting-state functional connectivity identifies individuals and predicts age in 8-to-26-month-olds. Developmental Cognitive Neuroscience, 56, 101123. 10.1016/j.dcn.2022.101123

Kleinhans, N. M., Müller, R.-A., Cohen, D. N., & Courchesne, E. (2008). Atypical functional lateralization of language in autism spectrum disorders. Brain Research, 1221, 115–125. 10.1016/j.brainres.2008.04.080

Klöppel, S., Mangin, J.-F., Vongerichten, A., Frackowiak, R. S. J., & Siebner, H. R. (2010). Nurture versus Nature: Long-Term Impact of Forced Right-Handedness on Structure of Pericentral Cortex and Basal Ganglia. The Journal of Neuroscience, 30(9), 3271–3275. 10.1523/JNEUROSCI.4394-09.2010

Klöppel, S., Vongerichten, A., Eimeren, T. van, Frackowiak, R. S. J., & Siebner, H. R. (2007). Can Left-Handedness be Switched? Insights from an Early Switch of Handwriting. Journal of Neuroscience, 27(29), 7847–7853. 10.1523/JNEUROSCI.1299-07.2007

Knecht, S., Dräger, B., Deppe, M., Bobe, L., Lohmann, H., Flöel, A., Ringelstein, E.-B., & Henningsen, H. (2000). Handedness and hemispheric language dominance in healthy humans. Brain, 123(12), 2512–2518. 10.1093/brain/123.12.2512

Kong, R., Spreng, R. N., Xue, A., Betzel, R. F., Cohen, J. R., Damoiseaux, J. S., De Brigard, F., Eickhoff, S. B., Fornito, A., Gratton, C., Gordon, E. M., Holmes, A. J., Laird, A. R., Larson-Prior, L., Nickerson, L. D., Pinho, A. L., Razi, A., Sadaghiani, S., Shine, J. M.,…Uddin, L. Q. (2025). A network correspondence toolbox for quantitative evaluation of novel neuroimaging results. Nature Communications, 16(1), 2930. 10.1038/s41467-025-58176-9

Kushner, H. I. (2012). Retraining left-handers and the aetiology of stuttering: The rise and fall of an intriguing theory. Laterality, 17(6), 673–693. 10.1080/1357650X.2011.615127

Kushner, H. I. (2013). Why are there (almost) no left-handers in China? Endeavour, 37(2), 71–81. 10.1016/j.endeavour.2012.12.003

Lebel, C., & Beaulieu, C. (2009). Lateralization of the arcuate fasciculus from childhood to adulthood and its relation to cognitive abilities in children. Human Brain Mapping, 30(11), 3563–3573. 10.1002/hbm.20779

Lee, M. H., Hacker, C. D., Snyder, A. Z., Corbetta, M., Zhang, D., Leuthardt, E. C., & Shimony, J. S. (2012). Clustering of Resting State Networks. PLoS ONE, 7(7), e40370. 10.1371/journal.pone.0040370

Lemaître, G., Nogueira, F., & Aridas, C. K. (2017). Imbalanced-learn: A Python Toolbox to Tackle the Curse of Imbalanced Datasets in Machine Learning. Journal of Machine Learning Research, 18, 1–5.

Li, D., Li, T., Niu, Y., Xiang, J., Cao, R., Liu, B., Zhang, H., & Wang, B. (2019). Reduced hemispheric asymmetry of brain anatomical networks in attention deficit hyperactivity disorder. Brain Imaging and Behavior, 13(3), 669–684. 10.1007/s11682-018-9881-5

Li, M., Chen, H., Wang, J., Liu, F., Long, Z., Wang, Y., Iturria-Medina, Y., Zhang, J., Yu, C., & Chen, H. (2014). Handedness-and Hemisphere-Related Differences in Small-World Brain Networks: A Diffusion Tensor Imaging Tractography Study. Brain Connectivity, 4(2), 145–156. 10.1089/brain.2013.0211

Lipkin, B., Tuckute, G., Affourtit, J., Small, H., Mineroff, Z., Kean, H., Jouravlev, O., Rakocevic, L., Pritchett, B., Siegelman, M., Hoeflin, C., Pongos, A., Blank, I. A., Struhl, M. K., Ivanova, A., Shannon, S., Sathe, A., Hoffmann, M., Nieto-Castañón, A., & Fedorenko, E. (2022). Probabilistic atlas for the language network based on precision fMRI data from >800 individuals. Scientific Data, 9(1), Article 1. 10.1038/s41597-022-01645-3

Lochy, A., de Heering, A., & Rossion, B. (2019). The non-linear development of the right hemispheric specialization for human face perception. Neuropsychologia, The Biological Basis of Social Cognition During Development, 126, 10–19. 10.1016/j.neuropsychologia.2017.06.029

Marek, S., & Dosenbach, N. U. F. (2018). The frontoparietal network: Function, electrophysiology, and importance of individual precision mapping. Dialogues in Clinical Neuroscience, 20(2), 133–140. 10.31887/DCNS.2018.20.2/smarek

Marek, S., Tervo-Clemmens, B., Calabro, F. J., Montez, D. F., Kay, B. P., Hatoum, A. S., Donohue, M. R., Foran, W., Miller, R. L., Hendrickson, T. J., Malone, S. M., Kandala, S., Feczko, E., Miranda-Dominguez, O., Graham, A. M., Earl, E. A., Perrone, A. J., Cordova, M., Doyle, O.,…Dosenbach, N. U. F. (2022). Reproducible brain-wide association studies require thousands of individuals. Nature, 603(7902), 654–660. 10.1038/s41586-022-04492-9

Mehta, K., Salo, T., Madison, T. J., Adebimpe, A., Bassett, D. S., Bertolero, M., Cieslak, M., Covitz, S., Houghton, A., Keller, A. S., Lundquist, J. T., Luo, A., Miranda-Dominguez, O., Nelson, S. M., Shafiei, G., Shanmugan, S., Shinohara, R. T., Smyser, C. D., Sydnor, V. J.,…Satterthwaite, T. D. (2024). XCP-D: A robust pipeline for the post-processing of fMRI data. Imaging Neuroscience, 2, 1–26. 10.1162/imag_a_00257

Meunier, D., Stamatakis, E. A., & Tyler, L. K. (2014). Age-related functional reorganization, structural changes, and preserved cognition. Neurobiology of Aging, 35(1), 42–54. 10.1016/j.neurobiolaging.2013.07.003

Mineroff, Z., Blank, I. A., Mahowald, K., & Fedorenko, E. (2018). A robust dissociation among the language, multiple demand, and default mode networks: Evidence from inter-region correlations in effect size. Neuropsychologia, 119, 501–511. 10.1016/j.neuropsychologia.2018.09.011

Miranda-Dominguez, O., Mills, B. D., Carpenter, S. D., Grant, K. A., Kroenke, C. D., Nigg, J. T., & Fair, D. A. (2014). Connectotyping: Model Based Fingerprinting of the Functional Connectome. PLOS ONE, 9(11), e111048. 10.1371/journal.pone.0111048

Muggeo, V. M. R. (2003). Estimating regression models with unknown break-points. Statistics in Medicine, 22(19), 3055–3071. 10.1002/sim.1545

Muggeo, V. M. R. (2008). segmented: An R Package to Fit Regression Models with Broken-Line Relationships. R News, 8(1).

Nenert, R., Allendorfer, J. B., Martin, A. M., Banks, C., Vannest, J., Holland, S. K., & Szaflarski, J. P. (2017). Age-related language lateralization assessed by fMRI: The effects of sex and handedness. Brain Research, 1674, 20–35. 10.1016/j.brainres.2017.08.021

Ocklenburg, S., & Güntürkün, O. (2024). Spatial attention, neglect, and the right hemisphere. In The Lateralized Brain (pp. 211–239). Elsevier. 10.1016/B978-0-323-99737-9.00006-9

Ojala, M., & Garriga, G. C. (2009). Permutation Tests for Studying Classifier Performance. 2009 Ninth IEEE International Conference on Data Mining, 908–913. 10.1109/ICDM.2009.108

Oldfield, R. C. (1971). The assessment and analysis of handedness: The Edinburgh inventory. Neuropsychologia, 9(1), 97–113. 10.1016/0028-3932(71)90067-4

Olsen, L. R. (2024). groupdata2: Creating Groups from Data (Version 2.0.5) [R]. 10.32614/CRAN.package.groupdata2

Olulade, O. A., Seydell-Greenwald, A., Chambers, C. E., Turkeltaub, P. E., Dromerick, A. W., Berl, M. M., Gaillard, W. D., & Newport, E. L. (2020). The neural basis of language development: Changes in lateralization over age. Proceedings of the National Academy of Sciences, 117(38), 23477–23483. 10.1073/pnas.1905590117

Papadatou-Pastou, M., Ntolka, E., Schmitz, J., Martin, M., Munafò, M. R., Ocklenburg, S., & Paracchini, S. (2020). Human Handedness: A Meta-Analysis. Psychological Bulletin, 146(6), 481–524. 10.1037/bul0000229

Pedregosa, F., Varoquaux, G., Gramfort, A., Michel, V., Thirion, B., Grisel, O., Blondel, M., Prettenhofer, P., Weiss, R., Dubourg, V., Vanderplas, J., Passos, A., Cournapeau, D., Brucher, M., Perrot, M., & Duchesnay, É. (2011). Scikit-learn: Machine Learning in Python. Journal of Machine Learning Research, 12(85), 2825–2830. 10.1016/j.patcog.2011.04.006

Perez, D. C., Dworetsky, A., Braga, R. M., Beeman, M., & Gratton, C. (2023). Hemispheric Asymmetries of Individual Differences in Functional Connectivity. Journal of Cognitive Neuroscience, 35(2), 200–225. 10.1162/jocn_a_01945

Peters, M. (1992). How sensitive are handedness prevalence figures to differences in questionnaire classification procedures? Brain and Cognition, 18(2), 208–215. 10.1016/0278-2626(92)90079-2

Philip, B. A., & Frey, S. H. (2014). Compensatory Changes Accompanying Chronic Forced Use of the Nondominant Hand by Unilateral Amputees. Journal of Neuroscience, 34(10), 3622–3631. 10.1523/JNEUROSCI.3770-13.2014

Poeppel, D. (2003). The analysis of speech in different temporal integration windows: Cerebral lateralization as ‘asymmetric sampling in time.’ *Speech Communication*, The Nature of Speech Perception, 41(1), 245–255. 10.1016/S0167-6393(02)00107-3

Power, J. D., Cohen, A. L., Nelson, S. M., Wig, G. S., Barnes, K. A., Church, J. A., Vogel, A. C., Laumann, T. O., Miezin, F. M., Schlaggar, B. L., & Petersen, S. E. (2011). Functional Network Organization of the Human Brain. Neuron, 72(4), 665–678. 10.1016/j.neuron.2011.09.006

Przybylski, L., & Kroliczak, G. (2023). The functional organization of skilled actions in the adextral and atypical brain. Neuropsychologia, 191, 108735. 10.1016/j.neuropsychologia.2023.108735

Rainio, O., Teuho, J., & Klén, R. (2024). Evaluation metrics and statistical tests for machine learning. Scientific Reports, 14(1), 6086. 10.1038/s41598-024-56706-x

Ratnarajah, N., Rifkin-Graboi, A., Fortier, M. V., Chong, Y. S., Kwek, K., Saw, S.-M., Godfrey, K. M., Gluckman, P. D., Meaney, M. J., & Qiu, A. (2013). Structural connectivity asymmetry in the neonatal brain. NeuroImage, 75, 187–194. 10.1016/j.neuroimage.2013.02.052

Salo, T., Mehta, K., Adebimpe, A., Bertolero, M., Murtha, K., Cieslak, M., Meisler, S., Madison, T., Sydnor, V., Covitz, S., Fair, D., & Satterthwaite, T. (2025). *XCP-D: A Robust Postprocessing Pipeline of fMRI data* (Version 0.10.5) [Computer software]. Zenodo. 10.5281/zenodo.14751221

Satterthwaite, T. D., Elliott, M. A., Gerraty, R. T., Ruparel, K., Loughead, J., Calkins, M. E., Eickhoff, S. B., Hakonarson, H., Gur, R. C., Gur, R. E., & Wolf, D. H. (2013). An improved framework for confound regression and filtering for control of motion artifact in the preprocessing of resting-state functional connectivity data. NeuroImage, 64, 240–256. 10.1016/j.neuroimage.2012.08.052

Schaefer, A., Kong, R., Gordon, E. M., Laumann, T. O., Zuo, X.-N., Holmes, A. J., Eickhoff, S. B., & Yeo, B. T. T. (2018). Local-Global Parcellation of the Human Cerebral Cortex from Intrinsic Functional Connectivity MRI. Cerebral Cortex, 28(9), 3095–3114. 10.1093/cercor/bhx179

Seitzman, B. A., Gratton, C., Laumann, T. O., Gordon, E. M., Adeyemo, B., Dworetsky, A., Kraus, B. T., Gilmore, A. W., Berg, J. J., Ortega, M., Nguyen, A., Greene, D. J., McDermott, K. B., Nelson, S. M., Lessov-Schlaggar, C. N., Schlaggar, B. L., Dosenbach, N. U. F., & Petersen, S. E. (2019). Trait-like variants in human functional brain networks. Proceedings of the National Academy of Sciences, 116(45), 22851–22861. 10.1073/pnas.1902932116

Sha, Z., Pepe, A., Schijven, D., Carrión-Castillo, A., Roe, J. M., Westerhausen, R., Joliot, M., Fisher, S. E., Crivello, F., & Francks, C. (2021). Handedness and its genetic influences are associated with structural asymmetries of the cerebral cortex in 31,864 individuals. Proceedings of the National Academy of Sciences, 118(47), e2113095118. 10.1073/pnas.2113095118

Sha, Z., Schijven, D., Carrion-Castillo, A., Joliot, M., Mazoyer, B., Fisher, S. E., Crivello, F., & Francks, C. (2021). The genetic architecture of structural left-right asymmetry of the human brain. Nature Human Behaviour. 10.1038/s41562-021-01069-w

Sheffield, J. M., Repovs, G., Harms, M. P., Carter, C. S., Gold, J. M., MacDonald, A. W., Daniel Ragland, J., Silverstein, S. M., Godwin, D., & Barch, D. M. (2015). Fronto-parietal and cingulo-opercular network integrity and cognition in health and schizophrenia. Neuropsychologia, 73, 82–93. 10.1016/j.neuropsychologia.2015.05.006

Siebner, H. R., Limmer, C., Peinemann, A., Drzezga, A., Bloem, B. R., Schwaiger, M., & Conrad, B. (2002). Long-Term Consequences of Switching Handedness: A Positron Emission Tomography Study on Handwriting in “Converted” Left-Handers. The Journal of Neuroscience, 22(7), 2816–2825. 10.1523/JNEUROSCI.22-07-02816.2002

Skeide, M. A., & Friederici, A. D. (2016). The ontogeny of the cortical language network. Nature Reviews Neuroscience, 17(5), Article 5. 10.1038/nrn.2016.23

Son, S., Miyata, J., Mori, Y., Isobe, M., Urayama, S.-I., Aso, T., Fukuyama, H., Murai, T., & Takahashi, H. (2017). Lateralization of intrinsic frontoparietal network connectivity and symptoms in schizophrenia. Psychiatry Research Neuroimaging, 260(101723001), 23–28. (rayyan-135758299). 10.1016/j.pscychresns.2016.12.007

Spironelli, C., Marino, M., Mantini, D., Montalti, R., Craven, A. R., Ersland, L., Angrilli, A., & Hugdahl, K. (2023). fMRI fluctuations within the language network are correlated with severity of hallucinatory symptoms in schizophrenia. Schizophrenia, 9(1), 75. 10.1038/s41537-023-00401-9

Steyerberg, E. W., Eijkemans, M. J. C., Harrell, F. E., & Habbema, J. D. F. (2000). Prognostic modelling with logistic regression analysis: A comparison of selection and estimation methods in small data sets. Statistics in Medicine, 19(8), 1059–1079. 10.1002/(SICI)1097-0258(20000430)19:8%3C1059::AID-SIM412%3E3.0.CO;2-0

Stone, S. P., Halligan, P. W., & Greenwood, R. J. (1993). The Incidence of Neglect Phenomena and Related Disorders in Patients with an Acute Right or Left Hemisphere Stroke. Age and Ageing, 22(1), 46–52. 10.1093/ageing/22.1.46

Strike, L. T., Hansell, N. K., Chuang, K.-H., Miller, J. L., de Zubicaray, G. I., Thompson, P. M., McMahon, K. L., & Wright, M. J. (2023). The Queensland Twin Adolescent Brain Project, a longitudinal study of adolescent brain development. Scientific Data, 10(1), 195. 10.1038/s41597-023-02038-w

Szaflarski, J. P., Binder, J. R., Possing, E. T., McKiernan, K. A., Ward, B. D., & Hammeke, T. A. (2002). Language lateralization in left-handed and ambidextrous people: fMRI data. Neurology, 59(2), 238–244. (rayyan-135759475).

Szaflarski, J. P., Holland, S. K., Schmithorst, V. J., & Byars, A. W. (2006). fMRI study of language lateralization in children and adults. Human Brain Mapping, 27(3), 202–212. 10.1002/hbm.20177

Szaflarski, J. P., Schmithorst, V. J., Altaye, M., Byars, A. W., Ret, J., Plante, E., & Holland, S. K. (2006). A longitudinal functional magnetic resonance imaging study of language development in children 5 to 11 years old. Annals of Neurology, 59(5), 796–807. 10.1002/ana.20817

Tuladhar, A. M., Snaphaan, L., Shumskaya, E., Rijpkema, M., Fernandez, G., Norris, D. G., & Leeuw, F.-E. de. (2013). Default Mode Network Connectivity in Stroke Patients. PLOS ONE, 8(6), e66556. 10.1371/journal.pone.0066556

Turkeltaub, P. E. (2019). A Taxonomy of Brain-Behavior Relationships After Stroke. Journal of Speech, Language, and Hearing Research, 62(11), 3907–3922. 10.1044/2019_JSLHR-L-RSNP-19-0032

Turkeltaub, P. E., Martin, K. C., Laks, A. B., & DeMarco, A. T. (2025). Right hemisphere language network plasticity in aphasia. Brain, awaf308. 10.1093/brain/awaf308

Van Essen, D. C., Smith, S. M., Barch, D. M., Behrens, T. E. J., Yacoub, E., & Ugurbil, K. (2013). The WU-Minn Human Connectome Project: An overview. NeuroImage, Mapping the Connectome, 80, 62–79. 10.1016/j.neuroimage.2013.05.041

Wang, D., Buckner, R. L., & Liu, H. (2014). Functional Specialization in the Human Brain Estimated By Intrinsic Hemispheric Interaction. Journal of Neuroscience, 34(37), 12341–12352. 10.1523/JNEUROSCI.0787-14.2014

Weber, K. A., Teplin, Z. M., Wager, T. D., Law, C. S. W., Prabhakar, N. K., Ashar, Y. K., Gilam, G., Banerjee, S., Delp, S. L., Glover, G. H., Hastie, T. J., & Mackey, S. (2022). Confounds in neuroimaging: A clear case of sex as a confound in brain-based prediction. Frontiers in Neurology, 13. 10.3389/fneur.2022.960760

Willems, R. M., Van der Haegen, L., Fisher, S. E., & Francks, C. (2014). On the other hand: Including left-handers in cognitive neuroscience and neurogenetics. Nature Reviews Neuroscience, 15(3), 193–201. 10.1038/nrn3679

Wilson, S. M., & Schneck, S. M. (2021). Neuroplasticity in Post-Stroke Aphasia: A Systematic Review and Meta-Analysis of Functional Imaging Studies of Reorganization of Language Processing. Neurobiology of Language, 2(1), 22–82. 10.1162/nol_a_00025

Yang, Q., & Wu, X. (2006). 10 challenging problems in data mining research. International Journal of Information Technology & Decision Making, 05(04), 597–604. 10.1142/S0219622006002258

Yeo, B. T. T., Krienen, F. M., Sepulcre, J., Sabuncu, M. R., Lashkari, D., Hollinshead, M., Roffman, J. L., Smoller, J. W., Zöllei, L., Polimeni, J. R., Fischl, B., Liu, H., & Buckner, R. L. (2011). The organization of the human cerebral cortex estimated by intrinsic functional connectivity. Journal of Neurophysiology, 106(3), 1125–1165. 10.1152/jn.00338.2011

Yilmaz, A. E., & Demirhan, H. (2023). Weighted kappa measures for ordinal multi-class classification performance. Applied Soft Computing, 134, 110020. 10.1016/j.asoc.2023.110020

Zhang, C., Dougherty, C. C., Baum, S. A., White, T., & Michael, A. M. (2018). Functional connectivity predicts gender: Evidence for gender differences in resting brain connectivity. Human Brain Mapping, 39(4), 1765–1776. 10.1002/hbm.23950

Zhang, Z. (2025). Network Abnormalities in Ischemic Stroke: A Meta-analysis of Resting-State Functional Connectivity. Brain Topography, 38(2), 19. 10.1007/s10548-024-01096-6

