## Supplemental Material for "Hemispheric Fingerprinting: Identifying left and right hemispheres using functional connectivity"

### Supplemental Material 1: Group balancing

#### *SM 1.1: Cross-validation*

In the HCP sample, 971 participants had complete resting-state (n = 935) or language task data (n = 963). We performed demographic balancing on all 971 for simplicity. Fold creation was done (1) keeping individuals from the same family in the same folds and (2) with respect to gender, age, and EHI. We used the `mlr3` (ver. 1.7.1; Lang et al., 2019) and `mlr3resampling` R packages (ver. 2026.5.19; Hocking et al., 2026) to do the balancing. Models were balanced on EHI, using gender and the four HCP-supplied age groups (22 – 25; 26 – 30; 31 – 35; 36+) as categorical balancing units.

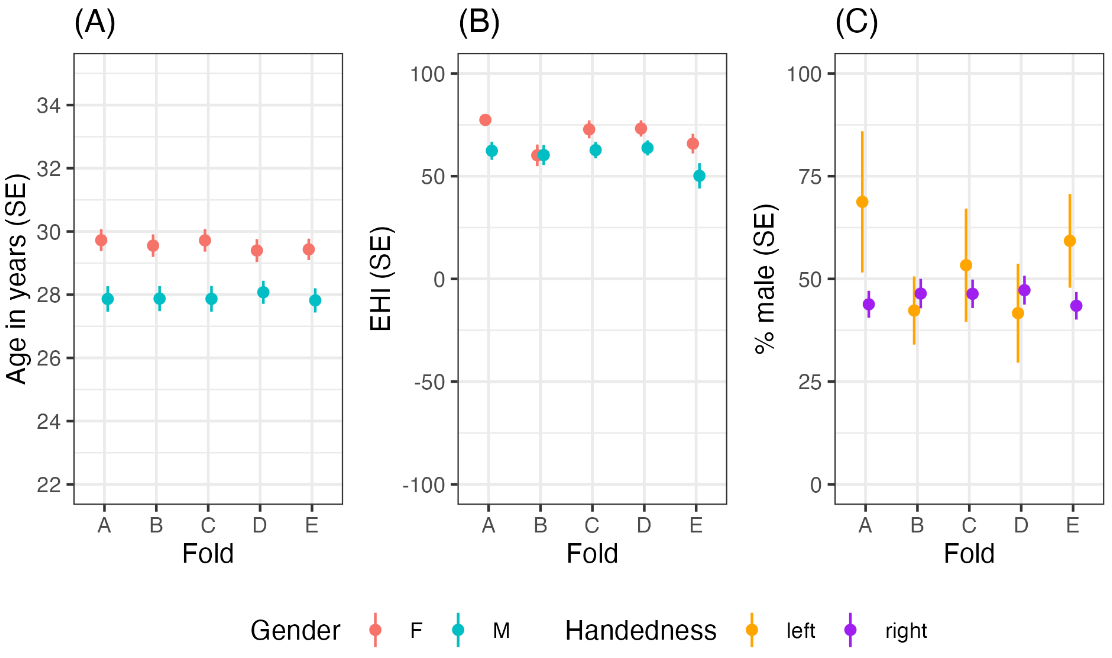


**Supplemental Figure 1.1:** (A) Average (SE) age in years by the five folds, divided by gender; (B) Mean (EHI) by fold, divided by gender; (C) Percent male (SE) by fold and handedness (i.e., EHI > 0: right-handed).

Supplemental Figure 1 shows the distribution of demographic variables across the five folds. As a sanity check, we ran a series of tests. A one-way ANOVA evaluating the effect of fold on exact age in years showed no differences between the groups, $F(4, 966) = 0.07, p = .99$. For EHI, which is not normally distributed, we used a Kruskal-Wallis test, which showed no effect of fold, $\chi^2 = 2.76, p = .60$. Third, we used a binomial GLM to test the different in percent-male between the folds and the `glht()` function from the R package `multcomp` (ver. 1.4-30; Hothorn et al., 2008) to show that there were no differences between groups (all $ps \geq .999$).

***SM 1.2: One hemisphere per participant***

Using the same parameters, participants were assigned to two LH/RH folds to balance tests run on one hemisphere at a time, see Supplemental Figure 2.

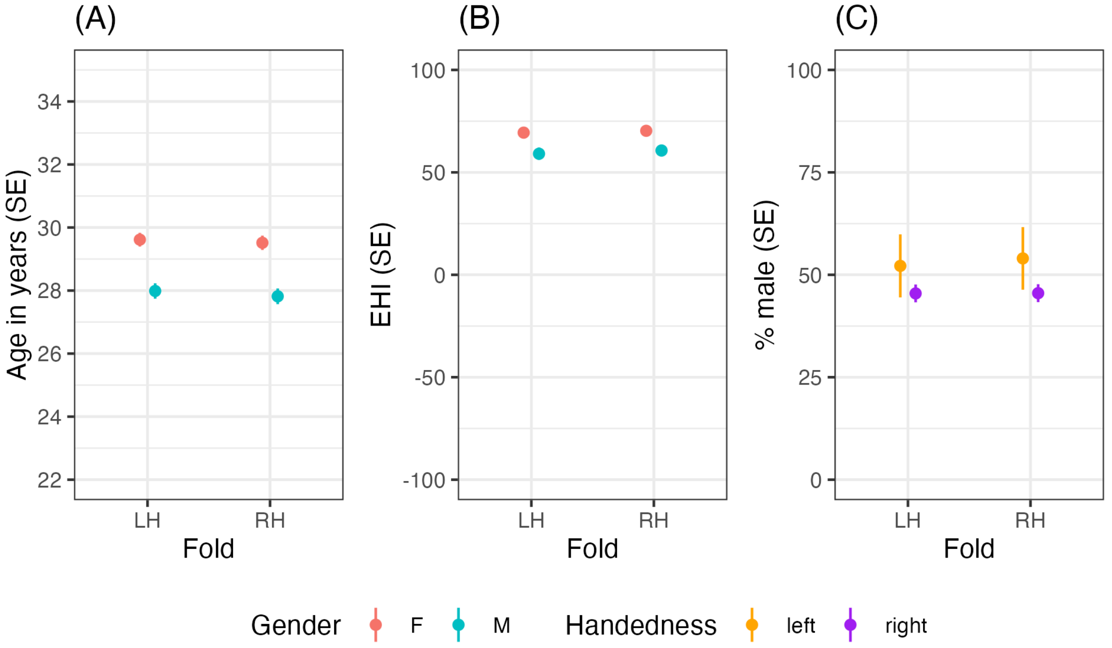


**Supplemental Figure 1.2:** (A) Average (SE) age in years by the two hemisphere folds, divided by gender; (B) Mean (EHI) by hemisphere fold, divided by gender; (C) Percent male (SE) by hemisphere fold and handedness (i.e., EHI > 0: right-handed).

A similar series of tests showed no differences between groups: age ANOVA: $F(1, 969) = 0.33$, $p = .57$; EHI Kruskal-Wallis: $\chi^2 = 1.21$, $p = .27$; and percent-male GLM, no effect of hemisphere (RH>LH): $\beta = 0.012$, $p = .925$.

***SM 1.3: Matching dextral samples***

In order to create similarly sized groups to compare with the lefties, we first randomly sample *n* dextrals from each fold, which were balanced demographically in SM 1.1 (*dextral-random*). In other words, all dextral individuals in Fold A were also in Fold A in all other mentions, etc. Next, using the R package `MatchIt` (ver. 4.7.3; Ho et al., 2011), we create demographically matched subsamples from each fold (*dextral-matched*). In this procedure, we balanced on absolute EHI value to account for the continuing imbalances in handedness strength.

As can be seen in Supplemental Figure 1.3.1(B), absolute EHI is more closely matched to the sinistrals (blue dots) in the dextral-matched group (red dots) than in the dextral-random group (green dots).

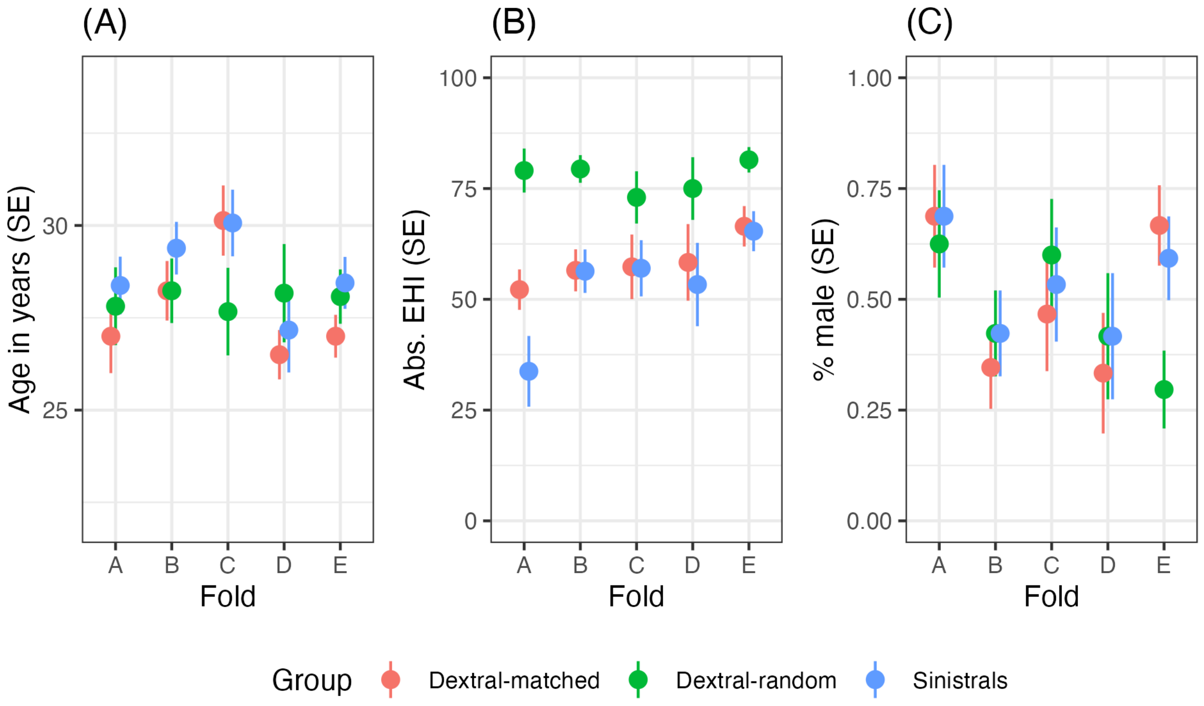


**Supplemental Figure 1.3:** (A) Average (SE) age in years in the sinistral and balanced dextral groups, by fold; (B) Mean absolute EHI by fold and handedness group; (C) Percent male (SE) by fold and handedness group.

In a similar series of models, a two-way ANOVA shows no effect of handedness group on mean age, $F(2, 281) = 1.86, p = .16$, nor an effect of fold, $F(4, 281) = 2.004, p = .09$. A Scheirer Ray Hare (R package `rcompanion` (ver. 2.5.2; Mangiafico, 2026)) test for two-way non-parametric comparisons for EHI shows an effect of handedness group, as expected, $H(2) = 44.9, p < .001$, and an effect of fold, $H(4) = 9.5, p = .05$. Finally, a binomial model shows that there is no effect of handedness groups (*p*s > .2), but there are fewer men in fold B relative to A (*β = -1.1, p = .003*) and in D relative to A (*β = -1.2, p = .012*).

The increasing number of constraints, especially the family structure in HCP make it difficult to create more balanced folds while also maximizing the number of participants (important for the non-dextral groups), thus we use these folds

**Supplemental Material 2: Network assignments**

Thirteen parcels received different labels in the LH and RH in the Cole-Antecevic parcellation. They are detailed in ST 2.1

**Supplementary Table 2.1:** ROIs receiving different labels in each hemisphere.

| **ROI** | **L Network** | **R Network** |
|---|---|---|
| PEF | Dorsal Attention | Cingulo Opercular |
| PSL | Language | Cingulo Opercular |
| STV | Language | Posterior Multimodal |
| 33pr | Cingulo Opercular | Frontoparietal |
| d32 | Default | Frontoparietal |
| 44 | Language | Frontoparietal |
| IFSp | Language | Frontoparietal |
| IFSa | Frontoparietal | Cingulo Opercular |
| OFC | Default | Frontoparietal |
| RI | Auditory | Somatomotor |
| STSda | Language | Default |
| 31a | Default | Frontoparietal |
| TE1m | Default | Frontoparietal |

**Supplementary Material 4: Preprocessing**

The automatically generated description of the XCP-D processing is shared below. This description was edited for clarity (additionally, see Salo, 2025).

> First, the motion parameters were calculated. The six translation and rotation head motion traces were low-pass filtered below 6.0 breaths-per-minute using a second-order Butterworth filter, based on Gratton et al. (2020). The Volterra expansion of these filtered motion parameters was then calculated. Framewise displacement (FD) was calculated from the filtered motion parameters (Power et al., 2014), with a head radius of 50 mm. Volumes with filtered FD values greater than 0.2 mm were flagged for the sake of later censoring (Power et al., 2014). In total, 27 nuisance regressors were used: the (#1 – 6) six motion parameters with their (#7 – 12) temporal derivatives, and (#13 – 24) the quadratic expansion of those 12 values; and (#25) mean global signal, (#26) mean white matter signal, and (#27) mean cerebrospinal fluid signal (Ciric et al., 2017; Satterthwaite et al., 2013). All motion parameters were filtered using the same parameters as described above and the Volterra expansion was calculated. The BOLD data were converted to NIfTI format, despiked with AFNI's 3dDespike (using the default settings, no mask, and the "NEW" option), and converted back to CIFTI format.
>
> Nuisance regressors were calculated from the BOLD data, replicating Nilearn's approach (Lindquist et al., 2019; Nilearn developers, 2024b; Salo, 2025), with the following exceptions. (1) Any volumes censored earlier in the workflow were first cubic-spline interpolated. (2) Outlier volumes at the beginning or end of the timeseries were replaced with the closest low-motion volume's values, to avoid extreme interpolations induced by cubic-spline interpolation. The resulting timeseries were band-pass filtered using a second-order Butterworth filter implemented in Nilearn (Nilearn developers, 2024a), in order to retain signals between 0.01 – 0.08 Hz. The same filter was applied to the confounds.
>
> After these steps, the band-pass-filtered timeseries were denoised using linear regression in Numpy (Harris et al., 2020). For this calculation, only the low-motion volumes were used to calculate parameter estimates. The interpolated volumes were denoised using the parameter estimates calculated from the low-motion volumes. The interpolated timeseries were then censored using the FD values calculated initially. See https://xcp-d.readthedocs.io/en/latest/workflows.html#denoising.
>
> Next, the denoised BOLD was then smoothed using Connectome Workbench (Marcus et al., 2011) with a Gaussian kernel (FWHM = 6.0 mm). Average within-region timecourses were extracted from the cleaned and filtered BOLD data using Connectome Workbench, using regions defined by the Glasser atlas. Pairwise functional connectivity (Pearson's r) between all regions was computed using Connectome Workbench. In cases of partial coverage,

uncovered vertices (values of all zeros or NaNs) were either ignored (when the parcel had > 50% coverage) or were set to zero (when the parcel had < 50% coverage).

Many internal operations of XCP-D use AFNI (Cox, 1996; Cox & Hyde, 1997), Connectome Workbench (Marcus et al., 2011), ANTs (Avants et al., 2009), TemplateFlow version 24.2.2 (Ciric et al., 2022), matplotlib version 3.10.0 (Hunter, 2007), Nibabel version 5.3.2 (Brett et al., 2022), Nilearn version 0.11.1 (Abraham et al., 2014), pybids version 0.18.1 (Yarkoni et al., 2019), and numpy version 2.2.1 and scipy version 1.15.1 (Harris et al., 2020). For more details, see the XCP-D website (https://xcp-d.readthedocs.io).

### Supplementary Material 4: MCC

In order to demonstrate the differences between accuracy and MCC, take the following example: There are 200 dextral (91%) and 20 sinistral (9%) individuals in our sample. As we iterate from 0/20 correctly predicted sinistrals to 20/20 correctly predicted sinistrals, the confusion matrix (ST 3.1) changes. For this example, the number of correctly predicted dextrals is held constant.

**Supplemental Table 4.1:** Example confusion matrices, for (left) 100% assignment for dextrals and changing sinistral success ratio and (right) vice versa for changing dextral success ratio.

| | | | | | | | | |
|---|---|---|---|---|---|---|---|---|
| Changing sinistral ratio | | | | | Changing dextral ratio | | | |
| | | Predicted class | | | | | Predicted class | |
| | | Sinistral | Dextral | | | | Sinistral | Dextral |
| True class | Sinistral | $x$ | *20-x* | | True class | Sinistral | 20 | 0 |
| | Dextral | 0 | 200 | | | Dextral | *200-x* | $x$ |

For “changing sinistral ratio,” as $x$ increases from 0 to 20, accuracy increases from $\frac{200}{220} = 0.91$ to 1.00, but MCC increases from 0 to 1. But, for example, a 50% error rate in the sinistral group leads to an MCC of 0.69, but an accuracy of 0.95.

For “changing dextral ratio,” as $x$ increases from 0, 200, accuracy increases from $\frac{20}{220} = 0.09$ to 1.00, and MCC increases from 0 to 1. However, as can be seen in Figure SM1.1, MCC is not completely insensitive to class size: A 50% error rate in the larger dextral group leads to an MCC of 0.29 (compared to $MCC = 0.69$ above), compared to an accuracy of 0.54.

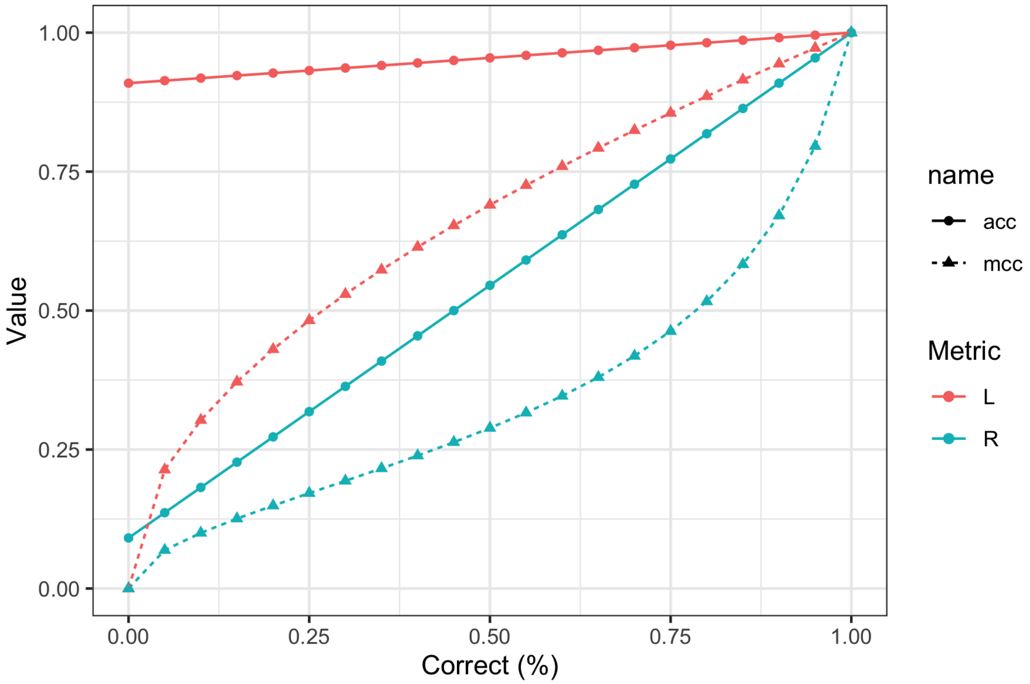


**Supplemental Figure 4.1:** Changes in accuracy (solid lines) and MCC (dashed lines) as the number of correctly predicted sinistrals (red) or dextrals (blue) changes, with the other group held constant at 100%.

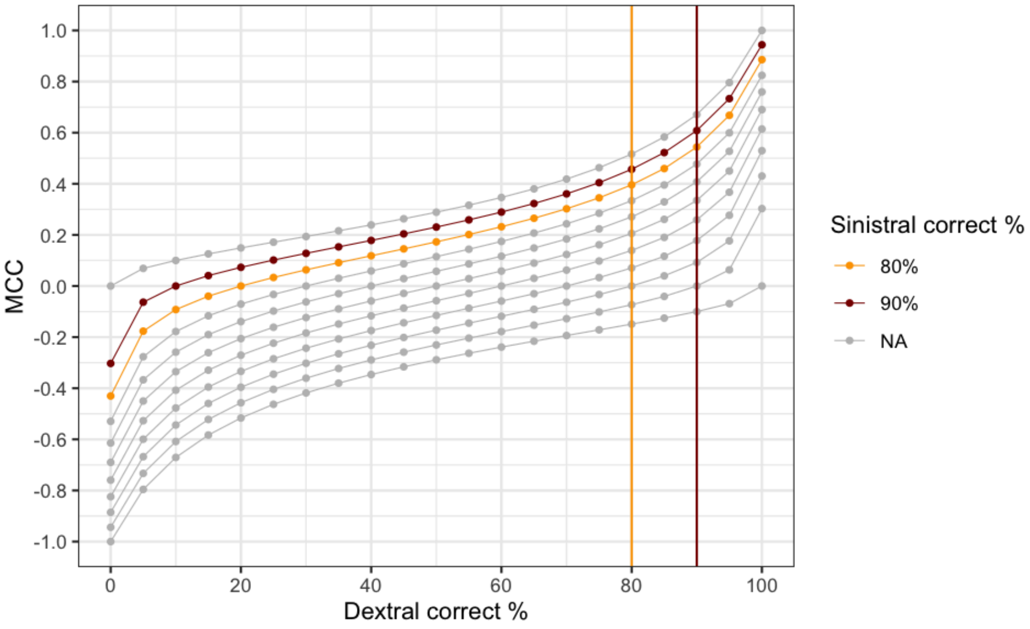


**Supplemental Figure 4.2:** Simulated MCC values (*y*-axis) for various values of dextral correct assignment rates (*x*-axis), by various values of sinistral correct assignment rates (lines). A correct assignment rate of 80% in each group is colored orange, and 90%, red.

**Supplemental Material 5: Results using neural networks and support vector classifiers**

We repeat the analyses described in the main text results using the neural net (NN) and support vector classifiers (SVC). Hemisphere classification performance for Analysis 1, repeated with NN and SVC are shown in ST 3.1.

**Supplemental Table 5.1:** Classifier performance (MCC) for neural networks (NN) and support vector classification (SVC) across data that include both hemispheres from each participant ("both") or one per participant ("1 per").

| **Input** | **Classifier** | **MCC (95% CI)** |
|---|---|---|
| Hemiconnectome (both) | NN | .991 [.985, .997] |
| Hemiconnectome (1 per) | | .983 [.971, .995] |
| Language task (both) | | .979 [.970, .988] |
| Language task (1 per) | | .962 [.944, .980] |
| Hemiconnectome (both) | SVC | .999 [.997, 1.0] |
| Hemiconnectome (1 per) | | 1.0 [1.0, 1.0] |
| Language task (both) | | .994 [.989, .999] |
| Language task (1 per) | | .985 [.974, .996] |

The examination of hemisphere, handedness, and overall classification performance is replicated with NN and SVC and shown in SF 3.1. Very similar patterns to the LDA classifier are evident. Namely handedness classification (red) fails across all tests, and hemisphere classification (green) remains nearly perfect in all tests.

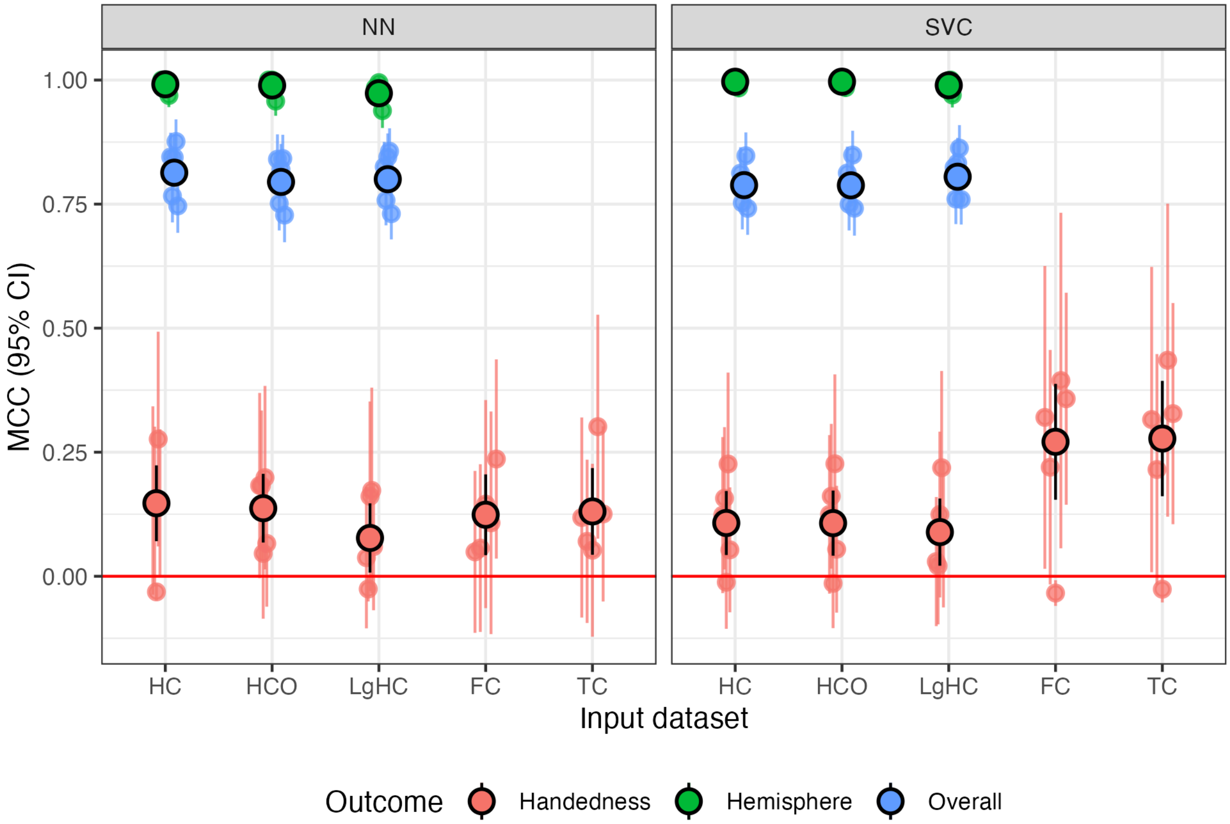


**Supplemental Figure 5.1:** MCCs for the handedness prediction across folds and input data using NN and SVC. The large, outlined points display the MCC calculated on the overall sample, and the smaller points, the MCC on a per-fold basis. HC: hemiconnectomes, HCO: hemiconnectomes oversampled; LgHC: language task hemiconnectomes; FC: full connectomes; TC: transconnectomes.

**Supplemental Material 6: Parcel asymmetries**

***SM 6.1: Parcel overlap***

Investigating the relationship between parcellations and asymmetries raises the question of whether asymmetries in the parcellation are driving the result. Other major parcellations are asymmetric, like the Glasser parcellation, including the Gordon-Laumann (Gordon et al., 2016) and Brainnetome atlases (Fan et al., 2016).

To index asymmetry, we calculated the Dice overlap between homotopes after projecting Glasser surface-space right-hemisphere ROIs onto the left hemisphere using Connectome workbench, where values closer to 0 indicate less symmetry. The Dice coefficients ranged from .45 (area LIPd) to .96 (area 3b). Homologues vary slightly in size, size $r = .97$.

Each connection therefore receives an combined asymmetry score of the average of the two Dice coefficients. The combined asymmetry score ranged from .46 (LIPd–STSva) to .96 (3b–V1). Based on the left-hemisphere Cole-Anticevic network assignments (Cole & Ito, 2018; Ji et al., 2019), the auditory and language networks were the most asymmetric (mean Dice: .69, .72) and the somatomotor and default mode networks were the most symmetric (mean Dice: .84, .82) out of the major networks.

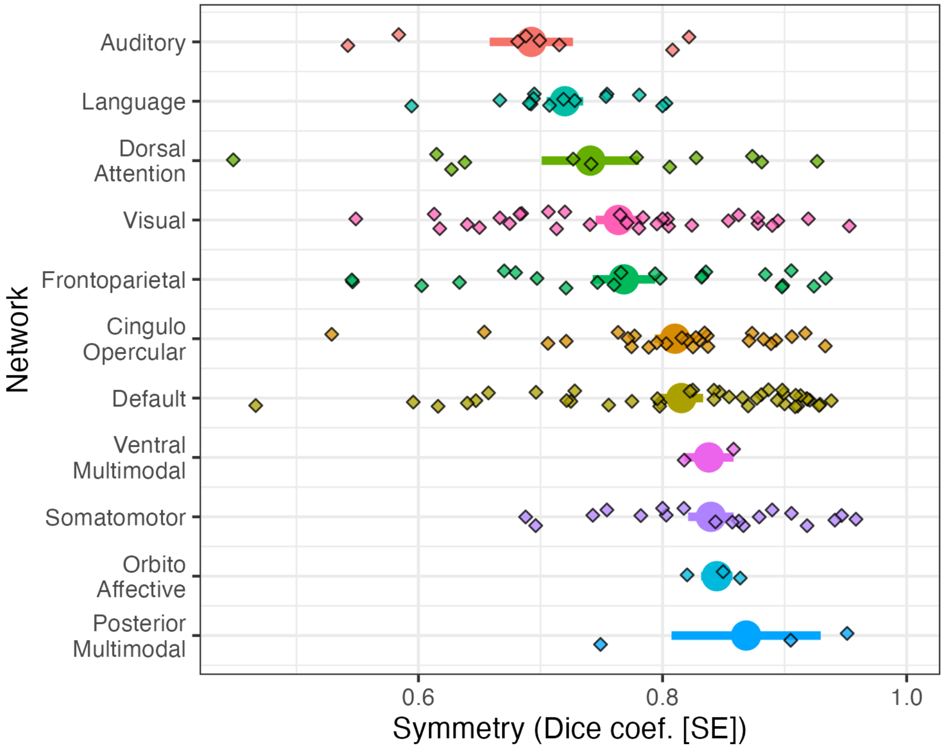


**Supplemental Figure 6.1:** Average (SE) Dice overlap for parcels by network. Individual regions are plotted as Xs. Higher numbers indicate greater symmetry with symmetry = 1 being perfect. The multimodal and orbito-affective networks all have three or fewer regions.

*SM 6.2: Relationship between asymmetry and feature importance*

As detailed in the main text, there is an association between *F* classifier score and symmetry of the parcels, but the effect is relatively minor, see Supplemental Figure 3.2.

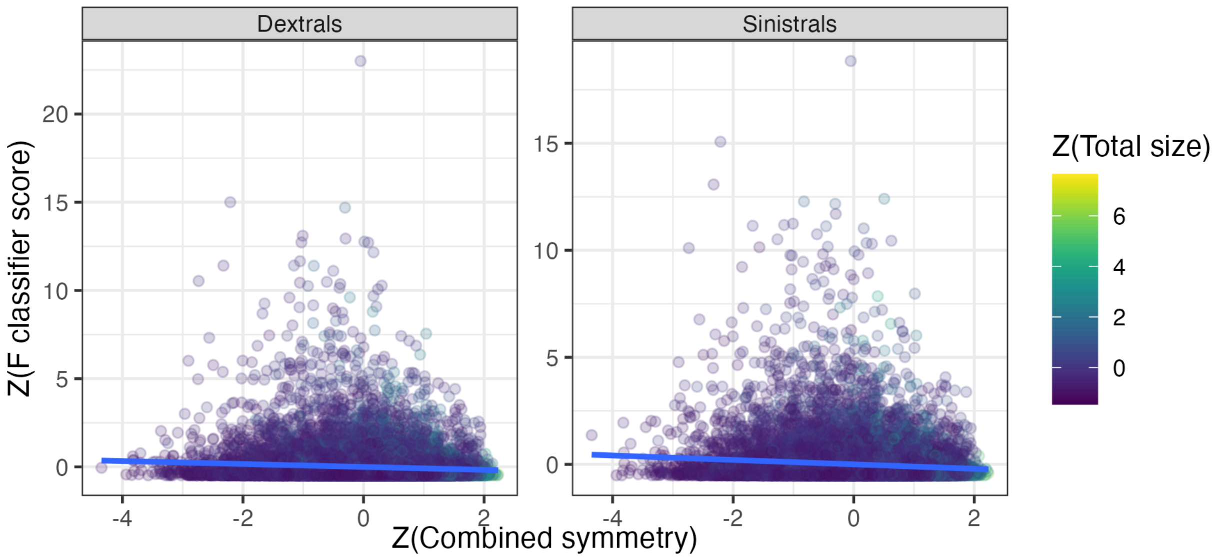


**Supplemental Figure 6.2:** *F* classifier score (scaled) vs. the combined symmetry score of the connection, colored by total size of the parcels involved in the connection.

**Supplemental Material 7: Enrichment sensitivity test**

This section contains the two enrichment sensitivity tests. We omit the per-connection plots for simplicity.

***SM 7.1: Excluded networks.***

First, we test permutations that re-include the three small, excluded networks, expanding the permutation space. No changes to the findings are found, see SF 6.1.

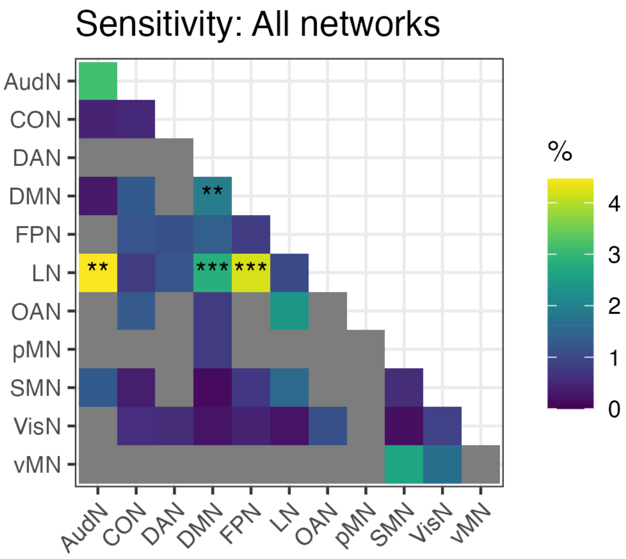


**Supplemental Figure 7.1:** Enrichment results including the OAN, vMN, and pMN. The same procedures are applied as in the main text.

***SM 7.2: Excluding parcels with different network assignments.***

Second, we exclude the ROIs that are part of different networks in each hemisphere, which tests that the networks identified as contributing to the differences between the hemispheres are not different only because they included some ROIs, but not others, in the RH. There were 13 of these parcels.

In this analysis, the LN-AudN connections were no longer significant. A new set of significant connections appears, CON-DMN, but this does not survive FDR correction, $p = .027$, $p_{FDR} = .246$

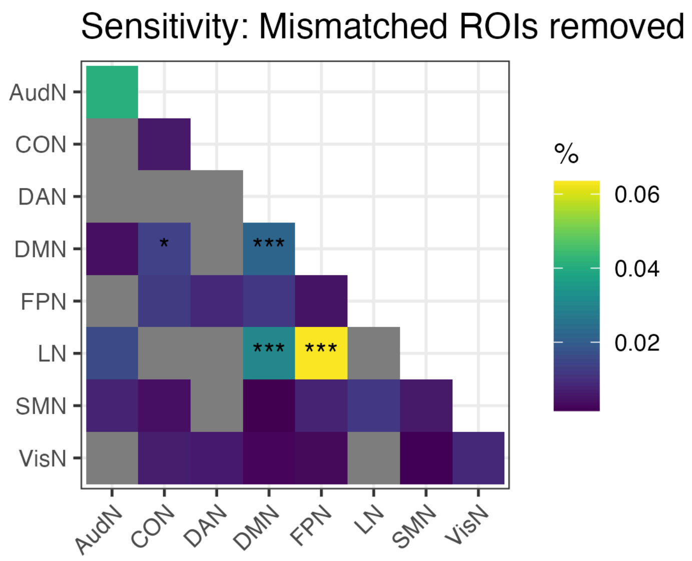


**Supplemental Figure 7.2:** Enrichment results after excluding 13 parcels that are in different networks in the two hemispheres. The same procedures are applied as in the main text.

The change in AudN-LN is likely due to the loss of several connections: one region that was in the AudN (RI), and five in the LN (PSL, STV, 44, IFSp, STSda). This nearly halves the already small total number of AudN-LN connections from $8 \times 14 = 112$ to $(8 - 1) \times (14 - 5) = 63$, which is the smallest set of connections outside of the within-AudN set (i.e., pre-removal: $8^2 = 64$, post-removal: $7^2 = 49$). This strains the limits of even bootstrapping for significance.

**Supplemental Material 8: Gender sensitivity test**

The main text describes that adding gender to the outcome did not improve classification success for handedness. If it did, we would expect the handedness MCC to increase going left-to-right (or from "No" to "Yes") in SF 4.1. Overall accuracy is improved due to the addition of high-accuracy gender cells to the classifier.

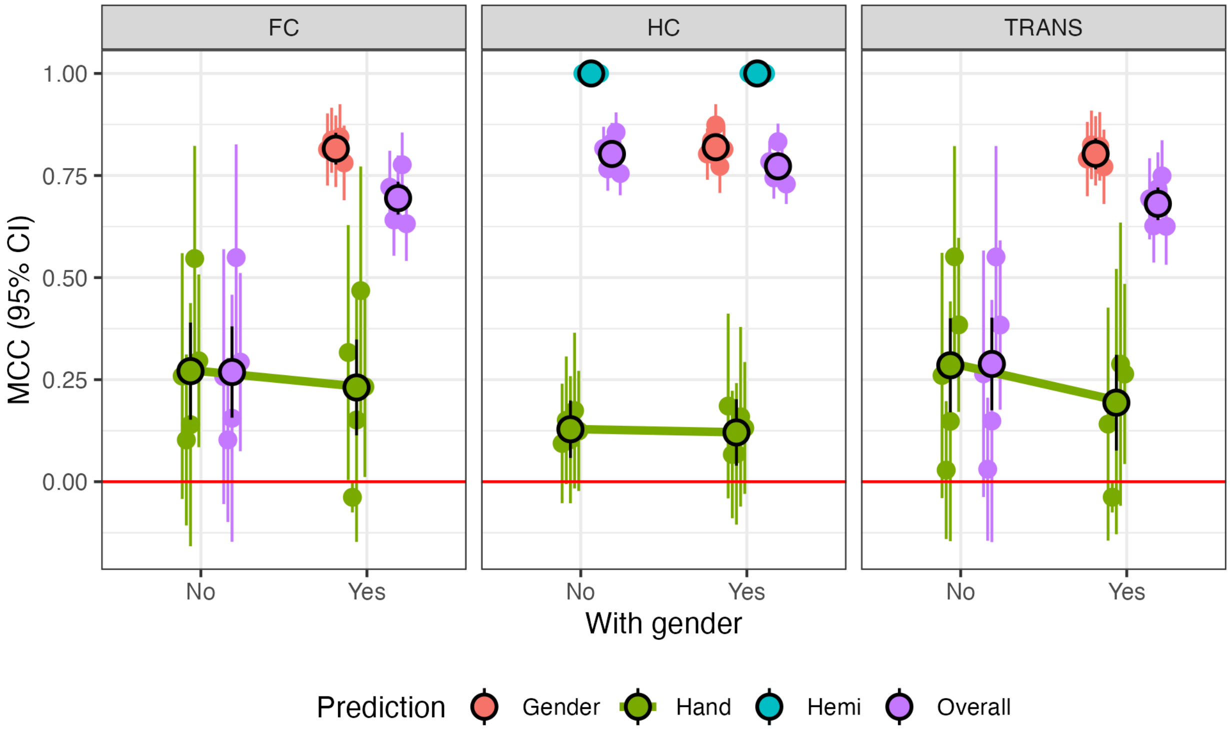


**Supplemental Figure 8.1:** Overall MCC (purple) as well as MCC for gender (red), handedness (green), and hemisphere (blue) without controlling for gender (left) and including gender (right). The green line between handedness points visually shows the change of interest. Naturally, the gender classification only appears in the "Yes" column.

### References


Abraham, A., Pedregosa, F., Eickenberg, M., Gervais, P., Mueller, A., Kossaifi, J., Gramfort, A., Thirion, B., & Varoquaux, G. (2014). Machine learning for neuroimaging with scikit-learn. *Frontiers in Neuroinformatics*, 14.

Avants, B. B., Tustison, N., & Song, G. (2009). Advanced normalization tools (ANTS). *Insight j*, *2*(365), 1–35.

Brett, M., Markiewicz, C. J., Hanke, M., Côté, M.-A., Cipollini, B., McCarthy, P., Jarecka, D., Cheng, C. P., Halchenko, Y. O., Cottaar, M., Larson, E., Ghosh, S., Wassermann, D., Gerhard, S., Lee, G. R., Wang, H.-T., Kastman, E., Kaczmarzyk, J., Guidotti, R., … freec84. (2022). *Nipy/nibabel:* (Version 4.0.0) [Computer software]. Zenodo. https://doi.org/10.5281/zenodo.591597

Ciric, R., Thompson, W. H., Lorenz, R., Goncalves, M., MacNicol, E., Markiewicz, C. J., Halchenko, Y. O., Ghosh, S. S., Gorgolewski, K. J., Poldrack, R. A., & others. (2022). TemplateFlow: FAIR-sharing of multi-scale, multi-species brain models. *bioRxiv*, 2021–02. https://doi.org/10.1101/2021.02.10.430678

Ciric, R., Wolf, D. H., Power, J. D., Roalf, D. R., Baum, G. L., Ruparel, K., Shinohara, R. T., Elliott, M. A., Eickhoff, S. B., Davatzikos, C., Gur, R. C., Gur, R. E., Bassett, D. S., & Satterthwaite, T. D. (2017). Benchmarking of participant-level confound regression strategies for the control of motion artifact in studies of functional connectivity. *NeuroImage, Cleaning up the fMRI Time Series: Mitigating Noise with Advanced Acquisition and Correction Strategies*, *154*, 174–187. https://doi.org/10.1016/j.neuroimage.2017.03.020

Cole, M. W., & Ito, T. (2018). *ColeLab/ColeAnticevicNetPartition: First public release of The Cole-Anticevic Brain-wide Network Partition (CAB-NP)* (Version v1.0.5-major) [Computer software]. Zenodo. https://doi.org/10.5281/zenodo.1455791

Cox, R. W. (1996). AFNI: Software for analysis and visualization of functional magnetic resonance neuroimages. *Computers and Biomedical Research, an International Journal*, *29*(3), 162–173.

Cox, R. W., & Hyde, J. S. (1997). Software tools for analysis and visualization of fMRI data. *NMR in Biomedicine: An International Journal Devoted to the Development and Application of Magnetic Resonance In Vivo*, *10*(4–5), 171–178.

Fan, L., Li, H., Zhuo, J., Zhang, Y., Wang, J., Chen, L., Yang, Z., Chu, C., Xie, S., Laird, A. R., Fox, P. T., Eickhoff, S. B., Yu, C., & Jiang, T. (2016). The Human Brainnetome Atlas: A New Brain Atlas Based on Connectional Architecture. *Cerebral Cortex*, *26*(8), 3508–3526. https://doi.org/10.1093/cercor/bhw157

Gordon, E. M., Laumann, T. O., Adeyemo, B., Huckins, J. F., Kelley, W. M., & Petersen, S. E. (2016). Generation and Evaluation of a Cortical Area Parcellation from Resting-State Correlations. *Cerebral Cortex*, *26*(1), 288–303. https://doi.org/10.1093/cercor/bhu239

Gratton, C., Dworetsky, A., Coalson, R. S., Adeyemo, B., Laumann, T. O., Wig, G. S., Kong, T. S., Gratton, G., Fabiani, M., Barch, D. M., Tranel, D., Miranda-Dominguez, O., Fair, D. A., Dosenbach, N. U. F., Snyder, A. Z., Perlmutter, J. S., Petersen, S. E., & Campbell, M. C. (2020). Removal of high frequency contamination from motion estimates in single-band fMRI saves data without biasing functional connectivity. *NeuroImage*, *217*, 116866. https://doi.org/10.1016/j.neuroimage.2020.116866

Harris, C. R., Millman, J. K., van der Walt, S. J., Gommers, R., Virtanen, P., Cournapeau, D., Wieser, E., Taylor, J., Berg, S., Smith, N. J., Kern, R., Picus, M., Hoyer, S., van Kerkwijk, M. H., Brett, M., Haldane, A., del Río, J. F., Wiebe, M., Peterson, P., … Oliphant, T. E. (2020). Array programming with NumPy. *Nature*, *585*(7825), 357–362. https://doi.org/10.1038/s41586-020-2649-2

Ho, D., Imai, K., King, G., & Stuart, E. A. (2011). MatchIt: Nonparametric Preprocessing for Parametric Causal Inference. *Journal of Statistical Software*, *42*, 1–28. https://doi.org/10.18637/jss.v042.i08

Hocking, T. D., Thibault, G., Bodine, C. S., Arellano, P. N., Shenkin, A. F., & Lindly, O. J. (2026). Same/Other/All K-Fold Cross-Validation for Estimating Similarity of Patterns in Data Subsets. *Statistical Analysis and Data Mining: An ASA Data Science Journal*, *19*(1), e70055. https://doi.org/10.1002/sam.70055

Hothorn, T., Bretz, F., & Westfall, P. (2008). Simultaneous Inference in General Parametric Models. *Biometrical Journal*, *50*(3), 346–363.

Hunter, J. D. (2007). Matplotlib: A 2D graphics environment. *Computing in Science & Engineering*, *9*(03), 90–95.

Ji, J. L., Spronk, M., Kulkarni, K., Repovš, G., Anticevic, A., & Cole, M. W. (2019). Mapping the human brain's cortical-subcortical functional network organization. *NeuroImage*, *185*, 35–57. https://doi.org/10.1016/j.neuroimage.2018.10.006

Lang, M., Binder, M., Richter, J., Schratz, P., Pfisterer, F., Coors, S., Au, Q., Casalicchio, G., Kotthoff, L., & Bischl, B. (2019). mlr3: A modern object-oriented machine learning framework in R. *Journal of Open Source Software*, *4*(44), 1903. https://doi.org/10.21105/joss.01903

Lindquist, M. A., Geuter, S., Wager, T. D., & Caffo, B. S. (2019). Modular preprocessing pipelines can reintroduce artifacts into fMRI data. *Human Brain Mapping*, *40*(8), 2358–2376. https://doi.org/10.1002/hbm.24528

Mangiafico, S. (2026). *rcompanion: Functions to Support Extension Education Program Evaluation* (Version 2.5.2) [Computer software]. https://cran.r-project.org/web/packages/rcompanion/index.html

Marcus, D. S., Harwell, J., Olsen, T., Hodge, M., Glasser, M. F., Prior, F., Jenkinson, M., Laumann, T., Curtiss, S. W., & Van Essen, D. C. (2011). Informatics and data mining tools and strategies for the human connectome project. *Frontiers in Neuroinformatics*, *5*, 4. https://doi.org/10.3389/fninf.2011.00004

Nilearn developers. (2024a). *Nilearn.signal.butterworth* (Version 0.12) [Python]. https://nilearn.github.io/dev/modules/generated/nilearn.signal.butterworth.html

Nilearn developers. (2024b). *Nilearn.signal.clean* (Version 0.12) [Python]. https://nilearn.github.io/modules/generated/nilearn.signal.clean.html

Power, J. D., Mitra, A., Laumann, T. O., Snyder, A. Z., Schlaggar, B. L., & Petersen, S. E. (2014). Methods to detect, characterize, and remove motion artifact in resting state fMRI. *NeuroImage*, *84*, 320–341. https://doi.org/10.1016/j.neuroimage.2013.08.048

Salo, T. (2025, October 28). *XCP-D boilerplate clarifications—Software Support* [Online post]. Neurostars. https://neurostars.org/t/xcp-d-boilerplate-clarifications/34426/2

Satterthwaite, T. D., Elliott, M. A., Gerraty, R. T., Ruparel, K., Loughead, J., Calkins, M. E., Eickhoff, S. B., Hakonarson, H., Gur, R. C., Gur, R. E., & Wolf, D. H. (2013). An improved framework for confound regression and filtering for control of motion artifact in the preprocessing of resting-state functional connectivity data. *NeuroImage*, *64*, 240–256. https://doi.org/10.1016/j.neuroimage.2012.08.052

Yarkoni, T., Markiewicz, C. J., de la Vega, A., Gorgolewski, K. J., Salo, T., Halchenko, Y. O., McNamara, Q., DeStasio, K., Poline, J.-B., Petrov, D., & others. (2019). PyBIDS: Python tools for BIDS datasets. *Journal of Open Source Software*, *4*(40).
